# Simultaneous brain-wide single-cell recording resolves spatiotemporal memory architecture

**DOI:** 10.64898/2026.05.21.726120

**Authors:** Dongqing Shi, Yongjie Hou, Yixiao Yan, Tian-hao Zhang, William C. Joesten, Peng Liu, Yixuan Wang, Mehul Gautam, Lirong Zheng, Jormay Lim, Jonathan Gould, BumJin Ko, Xiaoman Niu, Mou-Chi Cheng, Jung-Chien Hsieh, Florian Levet, Dawen Cai, Anne Draelos, Denise J. Cai, Donglai Wei, Changyang Linghu

**Author notes:** These authors contributed equally to this work (Dongqing Shi, Yongjie Hou).

## Abstract

Many fundamental mammalian brain functions emerge from the coordinated activity of cells distributed across large, brain-wide networks. To understand these processes in healthy and diseased states, ideally one would simultaneously measure and analyze single-cell activity at the brain-wide scale, an enduring challenge for live-measurement approaches that often face an inherent tradeoff between spatial resolution and scale. Here, we present GLOBE (sin<u>G</u>le-cell spatiotempora<u>L</u> rec<u>O</u>rding <u>B</u>rain-wid<u>E</u>), a technology for brain-wide single-cell recording of cellular activity *in vivo* with spatiotemporal resolution, physiological sensitivity, and parallelization-accelerated readout. GLOBE leverages genetically encoded intracellular protein tape recorders and a high-throughput computational platform for integrated image and signal analysis. GLOBE records analog signal amplitudes across a continuous time axis, requires only standard light microscopy for *in situ* readout, and is compatible with expansion microscopy and RNA readouts. We applied GLOBE to simultaneously record transcriptional activity of the immediate early gene *Fos* in up to 219,703 neurons simultaneously across a single mouse brain over 5.5 continuous days, with a timestamp precision of 3.1–6.7 hours (median absolute error), a local recording density of 69–90% of neurons per imaging field of view, and a post-mortem imaging readout speed of 2.9 seconds per neuron on average. GLOBE resolves the brain-wide spatiotemporal structure of single-cell activity, revealing that *Fos* transcriptional dynamics associated with fear learning and memory retrieval are distributed across the brain with region-specific temporal heterogeneity, and that the variance of this structure scales down as the number of sampled cells increases. We envision GLOBE to have broad applications for dissecting and decoding physiological and pathological processes at the brain-wide scale.

## 1 Introduction

Many physiological and pathological processes in the brain are driven and governed by cell populations distributed throughout the brain^1–5^. For example, a single fear memory can recruit neuronal ensembles spanning multiple brain regions, including the hippocampus, amygdala, prefrontal cortex, cingulate cortex, and thalamus^5–10^. These ensembles have been shown to evolve dynamically over time^10,11^, with recent theoretical and experimental evidence suggesting that the neurons encoding a memory can drift across time and experience even as the memory itself remains stable, a phenomenon known as representational drift^12,13^. Analogous observations have been made, at least in part, in other types of memory^14–16^. Furthermore, simultaneous chemogenetic reactivation of *Fos*+ cell ensembles distributed across multiple brain regions has been shown to induce stronger memory retrieval than reactivation of subsets in single regions, suggesting a causal role for coordinated ensembles distributed across the brain for memory function^5^. Dissecting these processes in healthy and diseased states requires a spatiotemporally resolved methodology capable of recording cellular activity across the entire brain.

Tetracycline-inducible (Tet-On/Tet-Off) and targeted recombination in active populations (TRAP) systems are popular in brain research, providing fluorescent reporter labeling of activated cell populations at a chemically defined time point and, when combined with end-point immunohistochemistry (IHC), allow snapshots at two time points with either Tet-On/Tet-Off or TRAP, or at three discrete time points when Tet-On/Tet-Off, TRAP, and IHC are combined in the same brain^17–20^. More recently, split-HaloTag-based chemical labeling has expanded this toolkit by capturing snapshots of cellular activity at time points defined by distinct fluorescent dyes^21,22^. However, because these systems rely on distinct fluorescent reporters, chemical dyes, or detection methods for each time point, the intensities of cell labels are not directly comparable across the discrete time points. Consequently, it remains challenging to quantify whether a cell’s activity has increased (activation), decreased (inhibition), or is unchanged relative to a prior state, a desired capability for studying longitudinal cellular dynamics underlying brain functions such as learning and memory^23^. Temporally continuous and longitudinally intensity-resolved recordings would enable direct, quantitative comparisons of cellular activity levels across the time axis, providing complementary information to existing fluorescent labeling-based readouts.

The *Fos* promoter is widely used to label and manipulate neuronal activity in learning and memory research, driving various synthetic payloads in mammalian brains such as fluorescent reporters^20,24–31^, chemically-controlled transactivators^20,24,26–30,32–35^, and optogenetic and chemogenetic molecular tools^20,24,27–30,33–35^. Although *in vivo* imaging of *Fos* promoter-driven fluorescent reporters provides temporally continuous and longitudinally intensity-resolved monitoring of cell populations over time, achieving brain-wide coverage at single-cell resolution remains challenging due to the inherent tradeoff between spatial resolution and imaging scale and the limited light access into tissue^36^.

Here, we introduce GLOBE, an *in vivo* protein tape recording technology with spatiotemporal resolution, brain-wide scale, and single-cell precision. GLOBE utilizes 1POK-based self-assembling protein tape recorders to record gene regulation dynamics in living cells for scalable post-mortem brain-wide readout, which has been shown to retain normal cellular physiology and function *in vitro* and *in vivo* and preserve intact brain functions in behaving mice^37–39^. GLOBE is fully genetically encoded, provides temporally continuous and longitudinally intensity-resolved recordings to support causal inference, and is compatible with expansion microscopy and RNA readouts in the same sample. GLOBE is empowered by a high-throughput computational platform for integrated image and signal analysis, with readout time scaling down proportionally through parallelization of microscopy (2.9 seconds per neuron per microscope) and computing (1.0 seconds per neuron per GPU). We demonstrated the capability of the GLOBE system by recording brain-wide single-cell *Fos* promoter activity simultaneously across up to 219,703 neurons per mouse over 5.5 continuous days, with a timestamp precision of 3.1–6.7 hours (median absolute error) and a local recording density of 69–90% of neurons per imaging field of view. GLOBE resolves the spatiotemporal structure of brain-wide single-cell activity *in vivo*, revealing brain region-specific spatial distributions of temporal transcriptional dynamics that correlate to contextual fear memory formation and retrieval, and that the variance of this structure scales down as the number of sampled cells increases. We envision GLOBE to provide a versatile recording platform with broad applications for biology and disease research at the brain-wide scale.

## 2 Results

### 2.1 System design of GLOBE recording

We first established the system design and brain-wide gene delivery approach for GLOBE recording (**Fig.1**), leveraging our previous molecular strategies for *in vivo* protein tape recording^37,38^. The GLOBE system consists of three modular, genetically encoded components delivered to single cells through brain-wide adeno-associated virus (AAV)-mediated transduction. The first, a constitutively expressed “structural monomer”, self-assembles into an intracellular protein assembly (referred to as “tape” throughout this paper) that elongates bidirectionally over time, forming a continuous molecular scaffold as the recording substrate. The remaining two components, “signal monomer” and “timestamp monomer”, are expressed at rates modulated by cellular activity and user-defined chemical inputs, respectively, and incorporated into this growing tape to encode simultaneous records of cellular activity and timestamp signals over time. Each monomer is fused to a distinct epitope tag for post hoc immunostaining readout of their relative location along the tape, to provide a time-resolved measurement of transcriptional activities within individual cells (**Fig. 1a**).

**Fig. 1.**
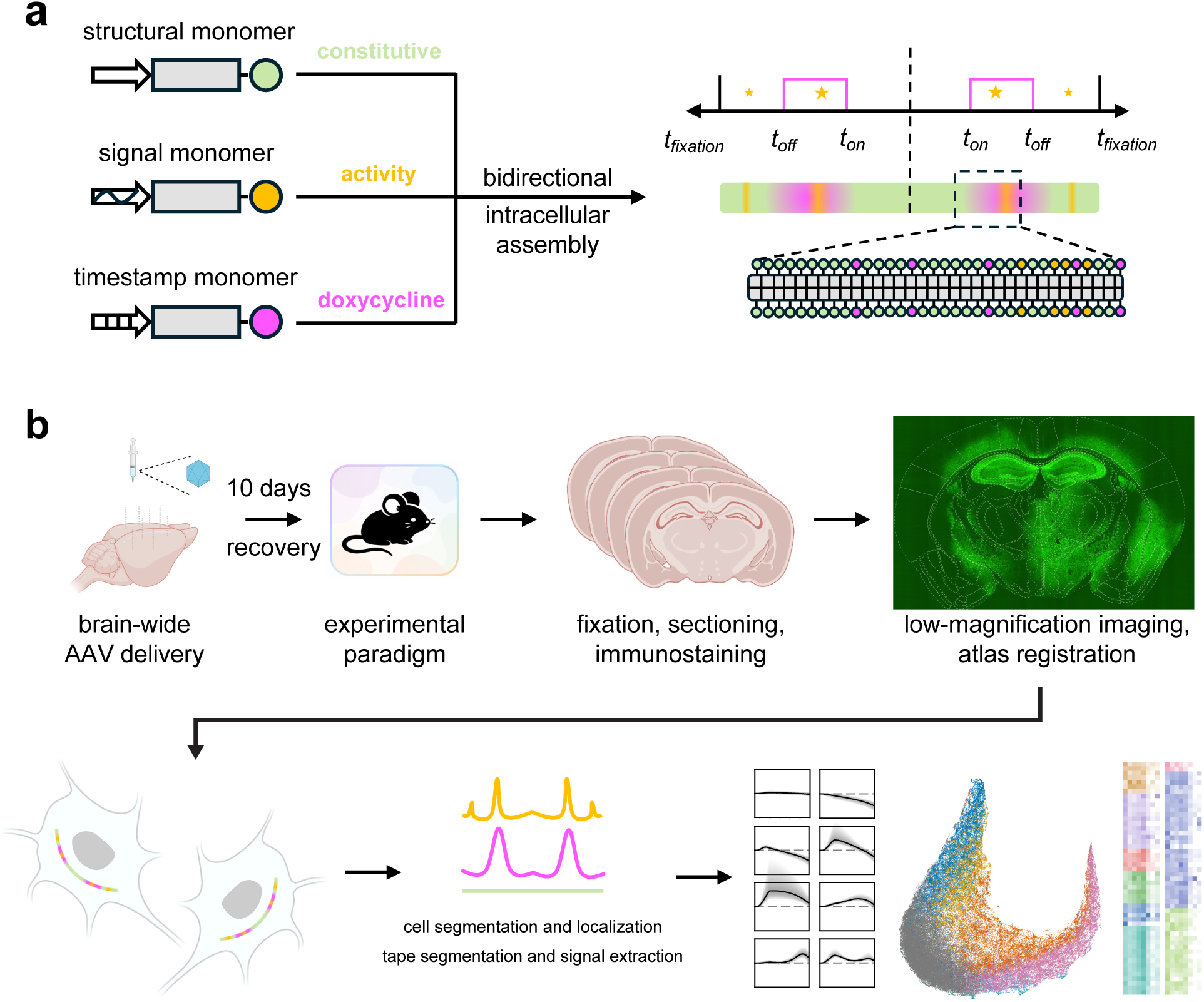
Design of the GLOBE recording system a, Molecular design of the GLOBE recording system. Constitutively expressed structural monomers, cell-activity-dependent signal monomers, and doxycycline (Dox)-dependent timestamp monomers co-assemble into bidirectionally elongating protein tapes in single cells. Temporal signals are encoded by distinct epitope tags on monomers, to be embedded in the spatial dimension along the tape for post-mortem imaging readout at scale. Stars denote experimental events**. b,** GLOBE workflow for brain-wide, single-cell, spatiotemporal activity recording, via brain-wide gene delivery, experimental paradigm (such as behavioral, pharmacological, or environmental events), immunohistochemistry, microscopy imaging, brain atlas registration, and high-throughput image analysis and spatiotemporal signal processing.

We set out to test whether brain-wide expression of the monomer of a protein tape recorder, XRI, which has been reported to enable days-long recording *in vivo*^37^, can be achieved. We performed systematic gene delivery via intravenous injection of AAV-PHP.eB or AAV.CAP-B10, two AAV serotypes with blood-brain barrier crossing capability, at the dosage of 1.7–3.4 x 10^11^ AAV genome copies (GC) per mouse, a comparable amount as used in literature^40,41^. This approach produced expression of structural monomer across the brain, yet in most brain regions only puncta or short tapes were observed in cells even 21 days after AAV delivery (**Extended Data Fig.1**), indicating insufficient expression levels for robust tape formation and sustained tape elongation for brain-wide recording. We then performed multi-site, multi-depth intracranial injections of AAV9 and observed broad expression coverage and robust tape formation in cells across the mouse brain (**Extended Data Fig. 2,3**). To evaluate whether this gene delivery strategy provides robust encoding of temporal transcriptional dynamics across the brain, we replicated our previously demonstrated *in vivo* recording paradigm in the hippocampus using the tamoxifen-inducible CreERT2 recombination^37^, while also expanding the recording coverage to additional brain regions within the same mouse. We observed robust recording of 4-OHT-induced transcriptional dynamics along tapes in cell populations across diverse brain regions *in vivo* (**Extended Data Fig.4**).

Having established the gene delivery approach, we next integrated the signal monomer and timestamp monomer modules into the system for GLOBE recording. We used the widely used mouse *Fos* promoter^32^ to drive the expression of V5-tagged XRI as the signal monomer, and TRE promoter-driven FLAG-tagged XRI as the timestamp monomer to be modulated by noninvasive, voluntary doxycycline (Dox) oral intake in a drinking water bottle in the home cage, to avoid potential confounding effects associated with injection-based administration, such as stress- or handling-induced *Fos* activity^42^. We performed multi-site, multi-depth AAV9 injections of the structural monomer, signal monomer, and timestamp monomer, together with rtTA3G driven by the human synapsin promoter across the mouse brain, at a total AAV dosage of 7.2 x 10^10^ GC per mouse (21%–42% of the per-mouse dosages used above for AAV-PHP.eB and AAV.CAP-B10). The AAV dosage ratio among the structural:signal:timestamp monomer constructs is set to 100:1:10, since we reasoned that a substantially larger proportion of the structural monomer relative to the signal and timestamp monomers would favor its dominant role in tape formation and elongation. We observed short tapes in the cell nucleus co-existing with tapes in the cytosol in 38% of neurons, and those nucleus-localized short tapes only contain the structural monomer but not the signal monomer or timestamp monomer (**Extended Data Fig.5**). We speculated that a small fraction of structural monomers, due to their higher expression level than those of signal and timestamp monomers, may have diffused into the cell nucleus in these cells before it had a chance to bind to the tape in the cytosol, resulting in the formation of nucleus-localized short tapes. Using the Nissl stain that selectively labels the cytosol of neuronal soma but not the cell nucleus, we computationally identified nucleus-localized short tapes and excluded them from further analysis throughout the rest of this paper.

### 2.2 Characterization and validation of GLOBE recording

To quantify the temporal accuracy and signal fidelity of GLOBE recording, we performed a time-series sampling experiment, collecting brains from animals sampled at defined time points along a shared experimental timeline (**Fig. 2a,b**). 10 days after hippocampal AAV delivery, Dox was administered in the drinking water (for the timestamp of Dox ON switch, defined as the onset of Dox-dependent signal rise). Dox was then withdrawn from the drinking water one day later (for the timestamp of Dox OFF switch, defined as the onset of Dox-dependent signal decay). The spatial encoding of temporal information along tapes were visualized from individual tapes acquired from brains collected at different time points, to resolve the temporal dynamics of Dox-dependent signal across mice (**Fig. 2c; Extended Data Fig. 6,7**). At each sampling time, we measured the fraction of tapes with Dox-dependent signal rise or decay, which provides an estimation of the cumulative latency distribution across tapes, where the median latency is defined by the half-maximal point of this cumulative distribution (**Fig. 2d,e**). Across mice, over 78% of all tapes successfully encoded the two timestamps on the final day of the timeline (15 days after AAV delivery). The median latency between the Dox administration and the appearance of the timestamp of Dox ON switch on the tape was 17.8 hours, and the median latency between the Dox withdrawal and the appearance of the timestamp of Dox OFF switch on the tape was 20.9 hours. We then aligned the two timestamps across all tapes to the real-world time axis by correcting for their respective median latencies. The remaining difference between this median latency-corrected recovered time and the real-world time was defined as the recovered time error, which reflects the inter-tape variations that cannot be globally corrected at the single-tape level, and thus provide an estimate of timestamp precision across individual tapes. The error for the recovered time had an IQR of -3.1 to 3.1 hours and -5.0 to 11.9 hours for the timestamps of Dox ON switch and OFF switch, respectively. Cumulative precision analysis showed the median absolute temporal errors are 3.09 and 6.69 hours for the timestamps of Dox ON switch and Dox OFF switch, respectively (**Fig. 2f,g**). These results indicate that GLOBE achieves hours-scale timestamp precision *in vivo*, enabling reconstruction of recorded activity along the real-world time axis.

**Fig. 2.**
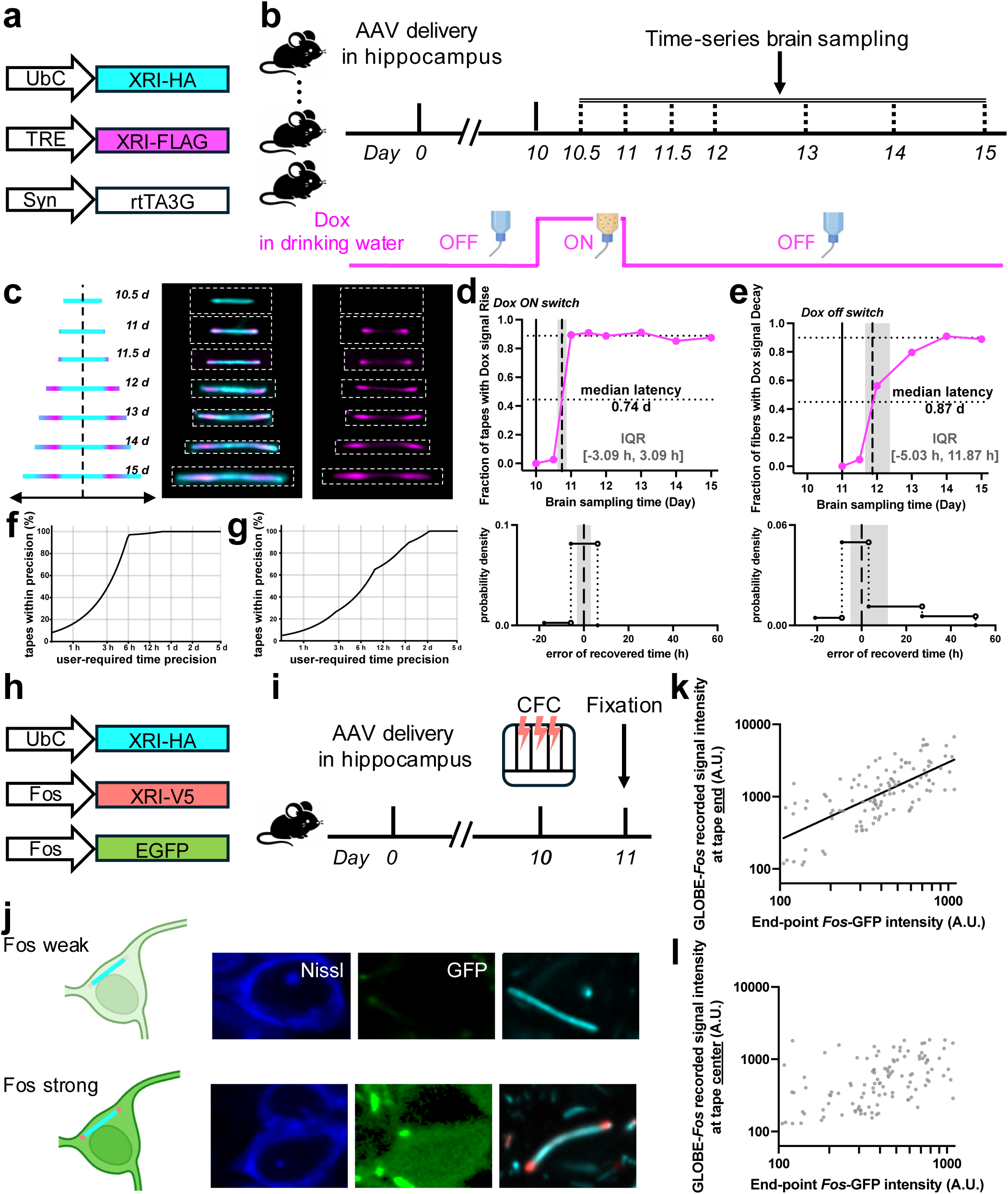
Characterization and validation of the GLOBE recording system in mice a, b, Construct schematics (**a**) and experimental timeline (**b**) for the characterization of GLOBE’s Dox-dependent timestamps via time-series brain sampling. AAVs were injected into the hippocampus on day 0, followed by Dox administration in drinking water from day 10 to day 11. Mice were perfused and brains were collected at defined times. **c,** Expected Dox-dependent timestamp signal distribution along the tapes (left) corresponds to representative confocal images (white dashed rectangles, boundaries) of single tapes stained for HA (cyan) and FLAG (Magenta) from brain sections collected at different sampling time points (middle, merged; right, FLAG only). See **Extended Data Fig. 6**,**7** for tissue-scale confocal images of cells and tapes. **d,** Quantification of Dox-dependent signal kinetics after the Dox ON switch. Top panel, fraction of tapes with Dox signal rise among all tapes across brain sampling times; bottom panel, error distribution of recovered time, defined as the difference between recovered time and real-world time after correcting for the median latency. Solid vertical line, Dox ON switch time; dotted horizontal lines, the plateau of tape fraction and the plateau-corrected half-maximum level for median latency calculation; dashed vertical line, time at median latency; shaded area, interquartile range (IQR) of the error of recovered time. *n* = 806 tapes from 7 brain sections from 2 mice at day 10.5, *n* = 320 tapes from 5 brain sections from 2 mice at day 11, *n* = 521 tapes from 3 brain sections from 1 mouse at day 11.5, *n* = 472 tapes from 4 brain sections from 2 mice at day 12, *n* = 526 tapes from 4 brain sections from 2 mice at day 13, *n* = 472 tapes from 3 brain sections from 3 mice at day 14, *n* = 312 tapes from 3 brain sections from 3 mice at day 15. **e,** Quantification of Dox-dependent signal kinetics after the Dox OFF switch, as in **d**. Top panel, fraction of tapes with Dox signal decay among all tapes with Dox signal rise. **f, g,** Fraction of tapes meeting user-defined temporal precision requirements, for Dox ON (**f**) or OFF (**g**) switch as the timestamp. **h, i,** Construct schematics (**h**) and experimental timeline (**i**) for cross-validation of *Fos* promoter activity recorded by GLOBE and by *Fos*-GFP under contextual fear conditioning (CFC), with results shown in **j**-**l**. AAVs were injected into the hippocampus on day 0, followed by CFC on day 10, and fixation on day 11, with immunostaining of HA (cyan) and V5 (Magenta). **j,** Schematic (left) and representative confocal images (right) of neurons with weak and strong *Fos* promoter activity as indicated by GFP fluorescence intensity. **k,** Correlation between GLOBE-recorded *Fos* signal intensity at the end of tape and endpoint *Fos*-GFP intensity. Each dot represents one neuron and one tape within. Black line, linear fit in log-log space. *n* = 114 neurons from 11 brain sections from 2 mice. **l,** Correlation between GLOBE-recorded *Fos* signal intensity at the center of tape and endpoint *Fos*-GFP intensity.

Since protein tape recorders embed the temporal axis along the spatial dimension, higher spatial resolution in imaging may improve the reconstruction accuracy of the time axis. We performed two Dox ON/OFF cycles on the mice and performed expansion microscopy^43,44^ for super-resolution imaging of the tapes in the mouse brain (**Extended Data Fig. 8a,b**). We observed expected timestamps sequentially aligned along single tapes, each corresponding to a Dox ON or OFF switch (**Extended Data Fig. 8c**). These results demonstrate that GLOBE is compatible with super-resolution microscopy, providing a potential framework for resolving temporal transcriptional events within single cells at higher resolution.

To validate that GLOBE faithfully reports transcriptional activity, we quantitatively compared GLOBE recording under *Fos* promoter-driven signal monomer with the labeling intensity of *Fos* promoter-driven GFP reporter (*Fos*-GFP, a well-established *Fos* transcriptional reporter^25,26,31,33^) in the same cells *in vivo*. Consistent with established literature^33^, AAV9-delivered *Fos*-GFP exhibited robust and condition-dependent induction across behavioral paradigms, including kainic acid-induced seizure and contextual fear conditioning, supporting the high efficiency of the AAV9-based gene delivery approach and the behaviorally relevant sensitivity of the *Fos*-promoter to couple *Fos* transcriptional activity to the expression level of synthetic payload *in vivo* (**Extended Data Fig. 9**). We then simultaneously applied GLOBE recording and *Fos*-GFP labeling, both delivered by AAV9, to the same hippocampal neuron populations *in vivo*, and fixed the brains 24 hours after contextual fear conditioning for imaging readout (**Fig. 2h,i**). In the same neurons, the intensity of *Fos*-GFP correlated with the amplitude of GLOBE-recorded signals at the time of fixation (at the tape ends). In contrast, GLOBE-recorded signals from an earlier time (at the tape center) in these neurons showed no visible correlation with *Fos*-GFP intensity (**Fig. 2j-l**). These results validate that GLOBE not only captures the real-world timing and temporal sequence of transcriptional events but also resolves the analog magnitude of behavior-associated transcriptional activity *in vivo*.

We further validated GLOBE’s compatibility with downstream cell-type and RNA readouts. We found that GLOBE-recorded samples are compatible not only with immunohistochemistry (IHC), as used for visualizing the tape monomers, but also with multi-round, spatially resolved RNA readout using RCA-based multiplexed RNA fluorescent *in situ* hybridization (FISH)^45^ (**Extended Data Fig. 10**; see **Methods** for the protocol). Such a combined single-cell spatial readout of protein tapes and RNAs in tissue may represent a promising approach to correlate GLOBE recordings to cell types and RNAs in the same cells, enabling a more comprehensive and integrative analysis of cellular processes across the brain, which is beyond the scope of this work on recording technology development.

### 2.3 GLOBE temporally resolves transcriptional dynamics associated with learning and memory

To test whether GLOBE can continuously resolve transcriptional activity associated with learning and memory, we applied the system to record *Fos* transcriptional dynamics in hippocampal neuron populations under contextual fear conditioning and recall (**Fig. 3a-c**). With the time axis reconstructed by Dox-dependent timestamps, we quantified *Fos* transcriptional dynamics during both the memory encoding and retrieval windows within the same animals. Mice that underwent contextual fear conditioning showed a significant increase in freezing during memory retrieval compared with the pre-conditioning period in the same context, confirming successful contextual fear memory encoding and retrieval (**Fig. 3d**). During both the encoding and retrieval windows, *Fos* transcriptional dynamics in experimental mice showed a significant distributional shift relative to time-matched home-cage control mice, which remained in the home cage throughout the experiment, with differences apparent in the upper tail of the distribution (**Fig. 3e,f**). These results demonstrate GLOBE’s ability to temporally resolve transcriptional activity with physiologically relevant sensitivity.

**Fig. 3.**
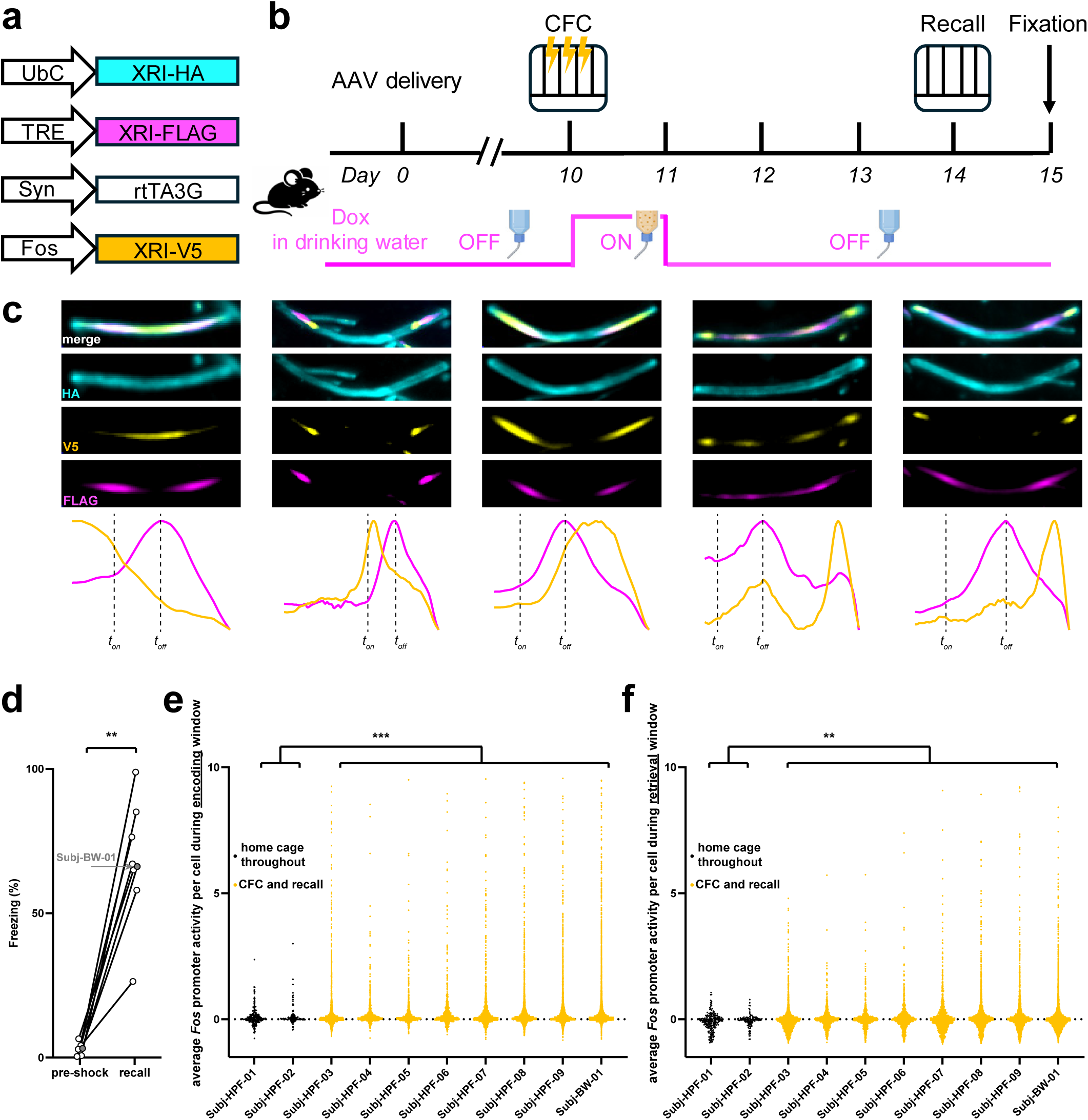
GLOBE captures single-cell *Fos* transcriptional dynamics during fear learning and memory retrieval a, b, Construct schematics (**a**) and experimental timeline (**b**) for GLOBE recording under contextual fear conditioning (CFC; day 10) and recall (day 14). **c,** Representative confocal images of tapes from hippocampal sections of mice experienced CFC and recall. Cyan, structural monomer via anti-HA immunostaining; yellow, Fos-promoter-driven monomer via anti-V5 immunostaining; magenta, Dox-dependent timestamp monomer via anti-FLAG immunostaining; blue, neuronal soma via Nissl staining. Fluorescence line profiles of V5 and FLAG channels in halves of the tapes are shown below, quantified by splitting each tape at its center, flipping the left half to align it with the right half, and averaging the two half-tape intensity profiles. **d,** Behavioral validation of contextual fear learning and memory retrieval. Freezing rate was measured during the 3 min pre-shock period on day 10 and then during the recall session on day 14 from the same mouse. Horizontal arrow, data from Subj-BW-01. **, P = 0.008, Wilcoxon matched-pairs signed rank test of freezing rate. *n* = 8 mice. **e,** GLOBE-recorded single-cell *Fos* transcriptional dynamics from the hippocampus, quantified as the average fold change during the contextual fear encoding window, defined as day 10 to day 11.5, relative to the baseline. Each dot represents one cell. Subj, mouse subject. Subj-HPF-01 to Subj-HPF-09 are mouse subjects with hippocampus-only AAV delivery, and Subj-BW-01 is a mouse subject with brain-wide AAV delivery and we analyzed here only the recordings from the hippocampus. Subj-HPF-01 and Subj-HPF-02 are home-cage controls (no CFC or recall). ***, P < 0.001, Anderson-Darling test of *Fos* promoter activity. *n* = 322 neurons from 2 mice for the home-cage control group, *n* = 19,904 neurons from 8 mice for the experiment group. **f,** Same as **c**, but for the retrieval window, defined as day 13.5 to day 15. **, P = 0.007, Anderson-Darling test of *Fos* promoter activity.

### 2.4 Brain-wide single-cell spatiotemporal recording

We next applied GLOBE for brain-wide, single-cell recording of *Fos* transcriptional dynamics over 5.5 continuous days, across contextual fear conditioning and then contextual fear recall 4 days later, resolving both the temporal activity and spatial location of individual cells (**Fig. 5a,b**). We developed an AI-powered, parallel-computing accelerated computational pipeline, GLOBE Tape Reader, for high-throughput brain-wide signal readout from tapes, cell segmentation, localization and registration to the Allen CCFv3 mouse brain atlas^46^, and recording quality check from microscopy images, building on our previous brain-region-specific computational framework, Tape Reader, for tape and cell segmentation^38^. Compared with Tape Reader, GLOBE Tape Reader employs an optimized neural network architecture retrained on new datasets containing human-annotated tape morphology in real microscopy images of tapes from diverse regions across the brain, with an average computational readout speed of 1.0 seconds per cell per GPU, (see **Methods** for details). GLOBE achieved broad coverage across major brain regions, including isocortex, olfactory area, cortical subplate, hippocampus, pallidum, striatum, thalamus, hypothalamus, and midbrain, recording up to 219,703 neurons simultaneously across a single mouse brain (**Fig. 4; Extended Data Fig. 11**).

**Fig. 4.**
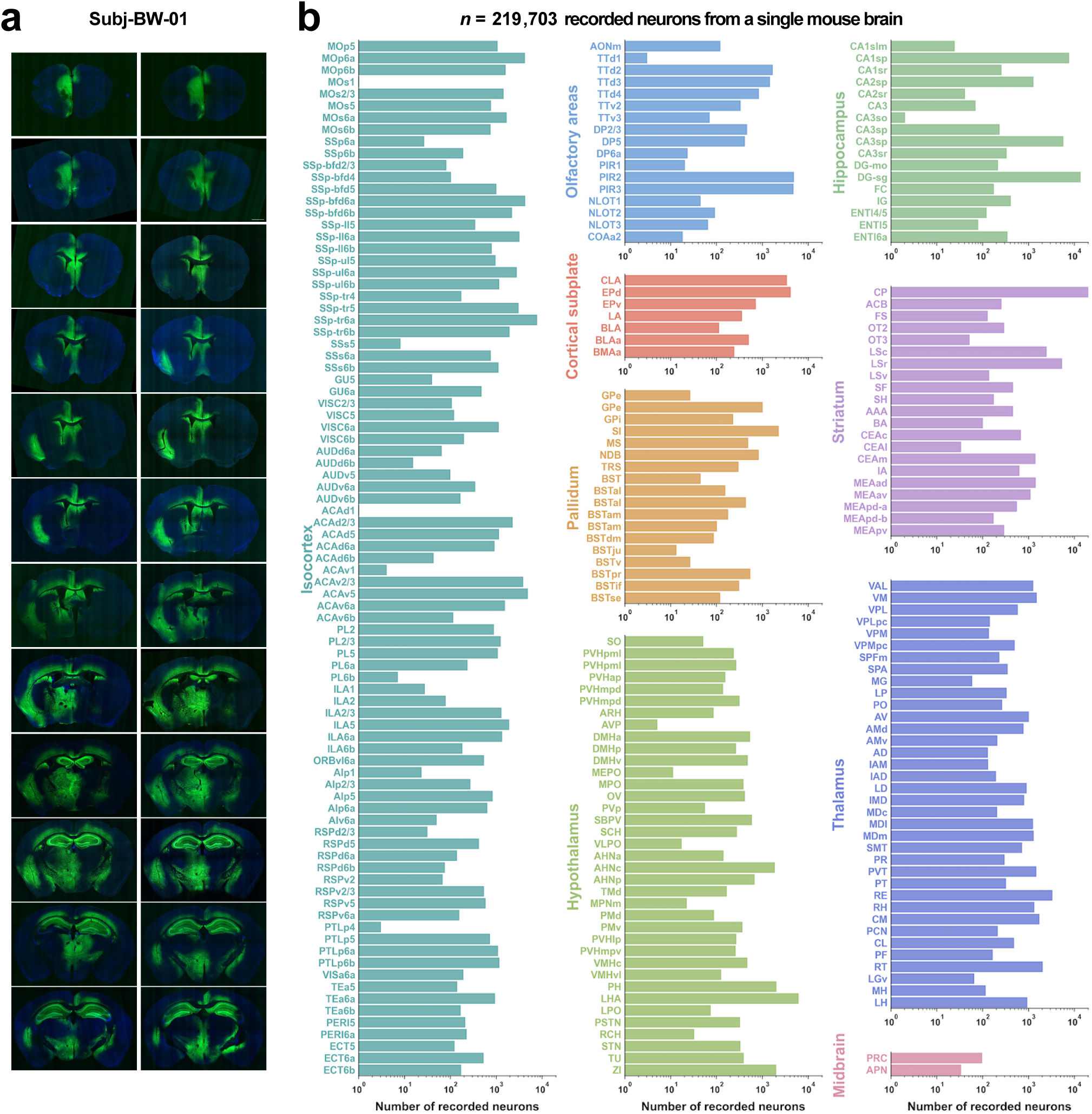
Brain-wide recording coverage in a GLOBE-recorded mouse a, Confocal images of coronal sections of a single mouse brain recorded by GLOBE under contextual fear conditioning and recall. Green, immunostaining for structural monomers; blue, Nissl stain for neurons. The brain was sectioned coronally at 30 µm thickness, with one section shown here per six consecutive sections. Sections are arranged from anterior to posterior (left-to-right and then top-to-bottom) at 0.18 mm intervals. See **Extended Data Fig. 2** for enlarged views. See **Fig. 5,6**,**7** for the full recording dataset and associated spatiotemporal analysis. **b,** Summary of the number of successfully recorded neurons, with Dox-dependent timestamps on tapes in them, across brain regions in the mouse shown in **a**, grouped by major anatomical divisions. *N* = 219,703 recorded neurons from 1 mouse.

**Fig. 5.**
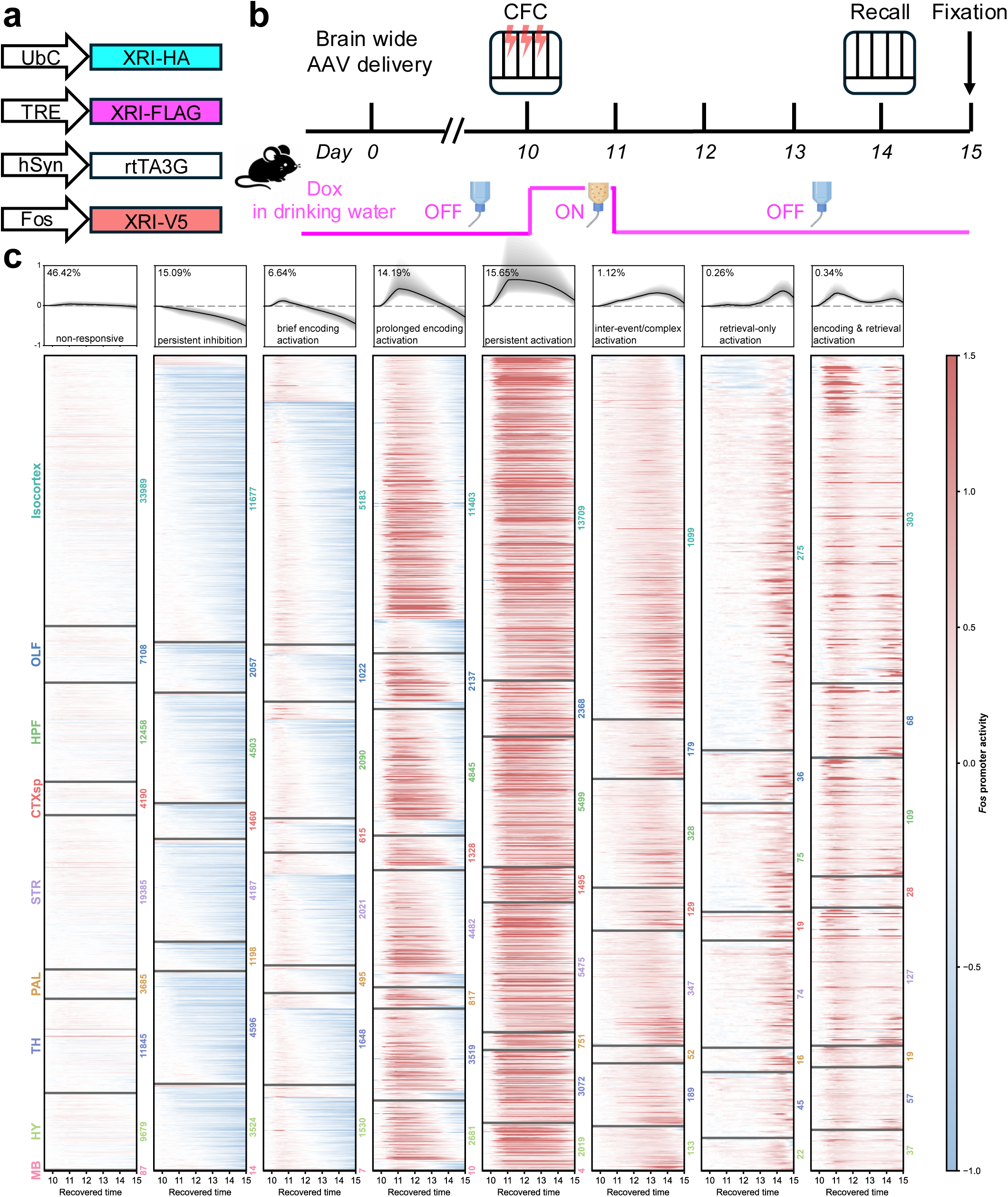
GLOBE reveals distinct temporal modes of brain-wide single-cell activity a, b, Construct schematics (**a**) and experimental timeline (**b**) for brain-wide GLOBE recording under contextual fear conditioning (CFC; day 10) and recall (day 14). **c,** GLOBE-recorded brain-wide single-cell *Fos* promoter activity between day 9.5 to day 15, quantified as the fold change relative to the baseline window (day 9.5 to day 10), grouped by eight distinct temporal modes. For each temporal mode: the top panel shows the median activity trajectory across all neurons in that mode, with the gray shading indicating nested percentile ranges of single-cell trajectories and the percentage indicating the abundance of neurons in that mode across all neurons; the bottom panel shows the recorded activity time course from single neurons (one neuron per row), with horizontal lines separating major anatomical divisions into sections and colored numbers indicating the number of neurons in each section. *n* = 219,703 recorded neurons from 1 mouse.

The recorded *Fos* transcriptional dynamics was then reconstructed along a continuous 5.5-day time axis spanning home-cage baseline, memory encoding, post-encoding period in the home cage, and memory retrieval. Machine-learning-based temporal trajectory classification identified eight distinct temporal modes of *Fos* transcriptional trajectories across neurons in the brain-wide recording dataset (**Fig. 5c**). The largest fraction of neurons (46%) was classified as the “non-responsive” mode, showing minimal *Fos* transcriptional modulation across the entire recording window.

Another group of neurons showed sustained mild suppression of *Fos* transcriptional activity after fear conditioning, a response pattern that is difficult to capture using conventional endpoint activity mapping methods, and were classified as the “persistent inhibition” mode. Among neurons with activity elevation after fear conditioning, some showed brief activity elevation and then decay on the day of memory encoding (the “brief encoding activation” mode), whereas others showed prolonged activity elevation lasting for 2–3 days (the “prolonged encoding activation” mode) or persistent activity elevation that continued through the retrieval period (the “persistent activation” mode)^47–49^. We also observed neurons with activity elevation emerging during the inter-event interval between encoding and retrieval or with waveforms more complex than simple activation or inhibition at the encoding or retrieval windows (the “inter-event/complex activation” mode). This observation suggests that activities in this temporal mode could arise from memory-associated processes after encoding, spontaneous activity unrelated to memory encoding or retrieval, or both, possibilities that could be further distinguished by future cellular perturbation of this population^10,13,23,50^. In addition, another group of neurons showed activity elevation only on the day of memory retrieval (the “retrieval-only activation” mode). Finally, a group of neurons showed two activity elevations, one on the day of encoding and another on the day of retrieval (the “encoding & retrieval activation” mode), indicating reactivation of encoding-associated neurons during memory retrieval. These temporal modes were distributed across diverse brain regions, in agreement with previous observations on the brain-wide distribution of memory-associated *Fos* activity^5^.

### 2.5 Spatiotemporal architecture of memory-associated brain-wide *Fos* transcriptional dynamics

We next investigated the spatiotemporal structure of the brain-wide recording dataset. We embedded the brain-wide single-cell *Fos* transcriptional dynamics by uniform manifold approximation and projection (UMAP) and examined how temporal features were organized across the manifold. By visualizing neurons by the overall magnitude and the temporal bias between encoding and retrieval window (referred to as the “encoding-retrieval index”) of their *Fos* transcriptional dynamics, we found that neurons with elevated *Fos* transcriptional dynamics formed a prominent arm on the right side of the manifold, while the temporal bias between encoding and retrieval windows formed a graded organization rather than a discretized organization across the manifold (**Fig. 6a,b**). The eight temporal modes identified above occupied distinct areas of the manifold (**Fig. 6c**). We further examined how these temporal modes were organized anatomically across space, and observed the relative abundance of the eight temporal modes varied substantially across brain regions (**Fig. 6d**), indicating that the *Fos* transcriptional dynamics associated with fear learning and memory retrieval are distributed at the brain-wide scale, with brain region-specific temporal heterogeneity.

**Fig. 6.**
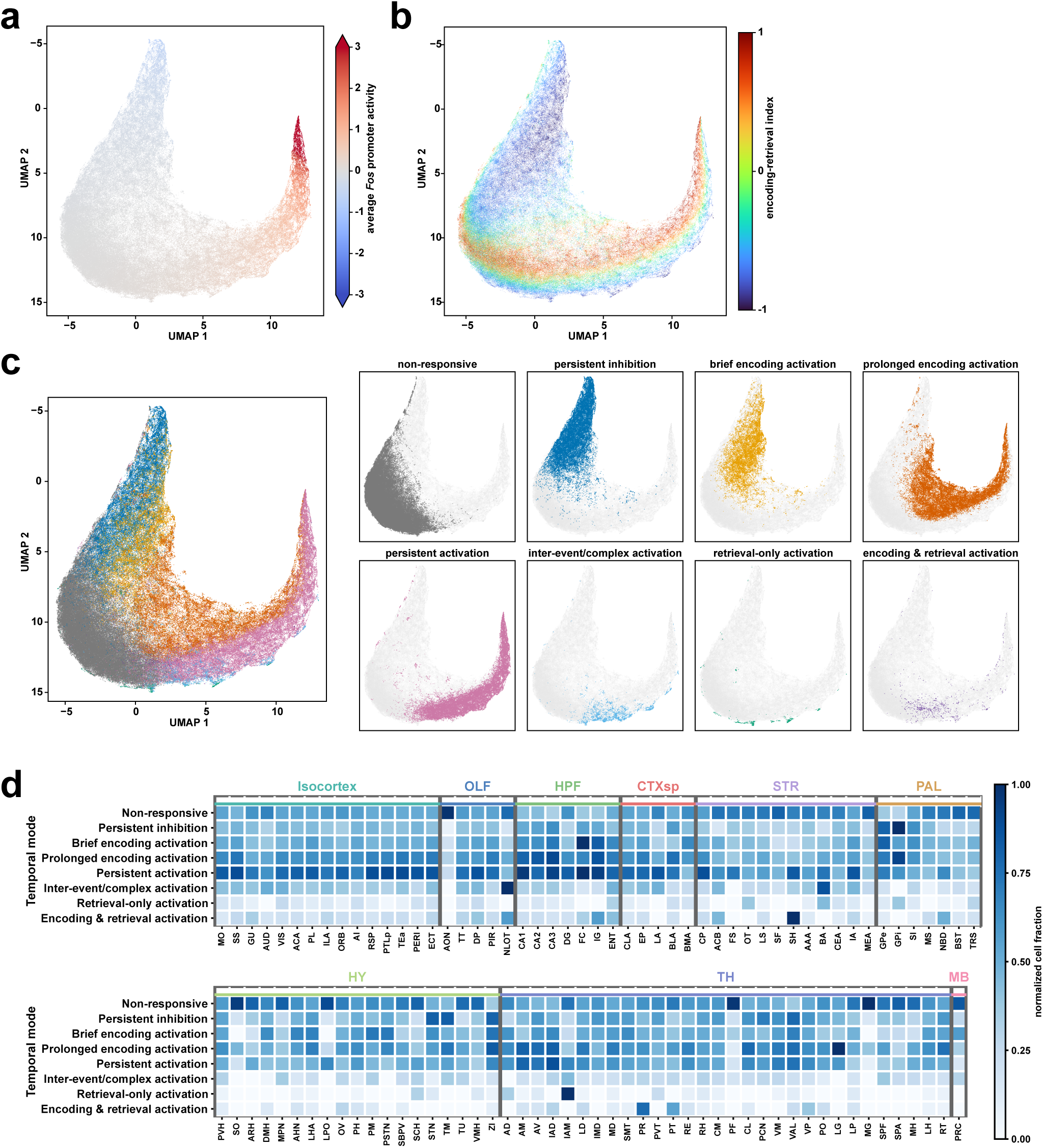
GLOBE reveals a brain-wide spatiotemporal atlas of single-cell activity from memory encoding through retrieval a, Uniform manifold approximation and projection (UMAP) temporal analysis of brain-wide single-cell *Fos* promoter activity. Each point represents one recorded neuron, colored by average *Fos* promoter activity across the full recording window. *n* = 219,703 recorded neurons from 1 mouse. **b,** As in **a**, but each neuron is instead colored by the encoding-retrieval index, quantifying the relative temporal bias of *Fos* promoter activity between encoding and retrieval windows. For each neuron, *Fos* promoter activity was averaged as absolute activity within the encoding-window, 10-11.5 d, and the retrieval-window, 13.5-15 d. The index was then calculated as the normalized difference between retrieval-window and encoding-window activity, encoding-window activity minus retrieval-window activity, divided by their sum. Positive values indicate stronger encoding-window activity, whereas negative values indicate stronger retrieval-window activity. **c,** Projection of classified neurons onto the UMAP space, colored by temporal mode. The right panels show the distributions of individual temporal modes on the UMAP. Gray points indicate the full recorded neuron population as a reference. **d,** Brain-region enrichment of distinct temporal modes. Color indicates the normalized cell fraction within each region, with darker color representing stronger enrichment of a given temporal mode. Brain regions with fewer than 50 recorded neurons were excluded from this analysis.

### 2.6 Population scaling refines the conserved manifold of brain-wide *Fos* transcriptional dynamics

We next investigated how population-level properties derived from the brain-wide recording dataset scale with the numbers of neurons within the same brain, up to the brain-wide scale. We examined the variance spectrum of principal component analysis (PCA) and its property under random cell subsampling. The full-population variance spectrum followed an approximate power-law decay with an exponent of 3.13, indicating that the majority of global temporal variance was captured by a small number of dominant, low-dimensional principal components (**Fig. 7a**). We randomly subsampled the recorded neuron population across various sample sizes and performed PCA for each sub-population, to compare their principal components (PCs) to those from the full-population. The dominant lowest-order PCs were highly stable even within small subsamples, while higher-order components required larger cell numbers to converge towards the full-population reference (**Fig. 7b**). Across repeated random cell subsampling, the variance of PC recovery decreased as the number of sampled cells increased (**Fig. 7c**). This observation provides evidence towards a memory coding principle that information coding is redundantly and widely distributed in cell populations across the brain, so that most information can be recovered from smaller subpopulations, while larger populations refine the precision of the information coding, resembling previous observations of sensory and behavioral information coding in the visual cortex via calcium imaging and electrophysiology^51–53^.

**Fig. 7.**
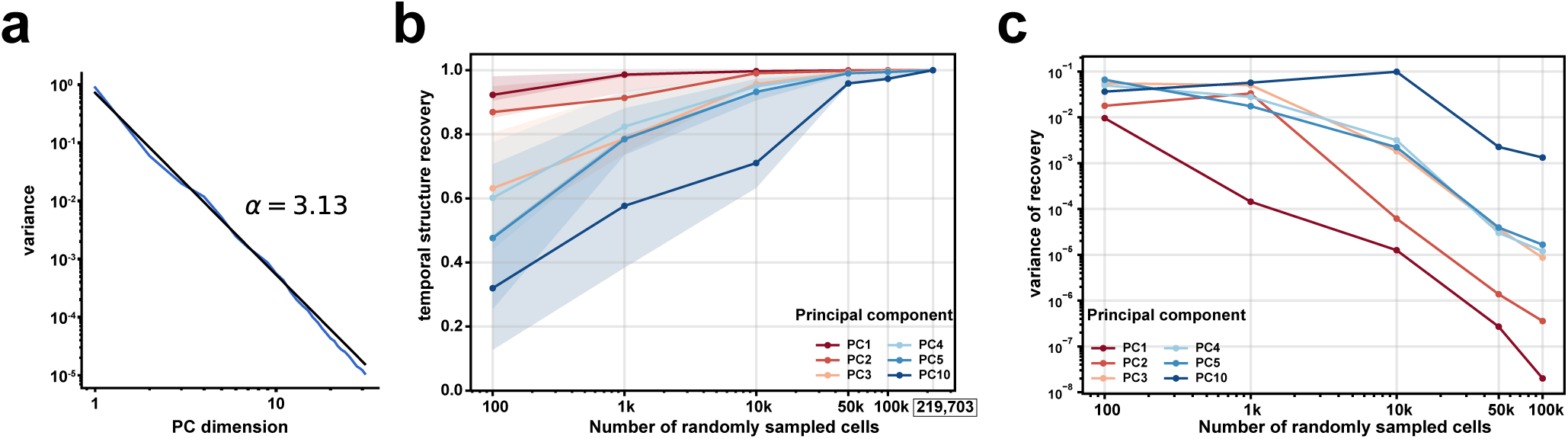
GLOBE reveals temporal structures of brain-wide single-cell activity a, Principal component (PC) variance spectrum (blue) of brain-wide single-cell *Fos* transcriptional dynamics. Black line, linear fit to the power-law form 1/*n*^α^, where *n* denotes the *n*-th PC dimension and *α* is the fitted power-law exponent (*α* = 3.13). **b,** Recovery of principal-component temporal structure across random cell subsampling. Cells were randomly subsampled at increasing population sizes, and PCA was recomputed independently for each subsample. The full population of 219,703 recorded neurons was used as the reference. Temporal structure recovery was quantified as the absolute Pearson correlation between the subsampled and full-population PC temporal loadings. Lines, the mean values across repeated random subsamplings; shaded boundaries, the interquartile ranges across repeats. 100, 50, 20, 20, 20 repeats were performed for subsample sizes of 100, 1k, 10k, 50k, 100k cells, respectively. **c**, For each pair of subsample size and PC, variance of recovery was computed across repeated random subsamplings. Lower variance of recovery indicates more reproducible recovery of the corresponding PC temporal feature across independent random subsamples.

## 3 Discussion

Simultaneous mammalian-brain-wide, single-cell recording of neural activity in the same subject remains a major unmet need in neuroscience^54,55^. In this work, we address this gap with genetically encoded protein tape recorders to enable brain-wide, single-cell recording of gene transcriptional activity. We found such recording provides physiological sensitivity *in vivo*, successfully resolving gene transcription activity induced by minutes-scale behavioral experiences, perhaps arising from the spontaneous molecular enrichment of reporter signals at the foci of tape termini. By recording and analyzing transcriptional correlates of memory encoding and retrieval across the brain, we provide a spatiotemporal activity atlas of memory as probed by the *Fos* promoter. We observed brain region-specific distributions of distinct temporal modes of *Fos* transcriptional dynamics, a gateway transcription factor involved in learning and memory, collectively forming a brain-wide activity manifold with its variance scales down as the number of sampled cells increases.

Building on our previous work establishing intracellular protein assemblies and proof-of-concept *in vivo* protein tape recording under chemically induced transcriptional activities^37,38^, GLOBE scales protein tape recording to the organ-wide regime and, via scaled and automated experimental and computational pipelines, demonstrates an unprecedented throughput—recording nearly a quarter-million spatially resolved neurons per mouse (**Extended Data Fig. 11**) over 5.5 continuous days at a local recording density of 69–90% of neurons per imaging field of view and an average readout speed of 2.9 seconds per neuron per microscope for image acquisition and 1.0 seconds per neuron per GPU for computational analysis. At this readout speed, imaging at the brain-wide scale reported here with one microscope followed by sequential single-GPU processing, the total readout time would be 10 days. This time scales down proportionally with parallelization, whether through multiple microscopes scanning sectioned tissues in parallel, multiple GPUs processing concurrently (as employed here), or processing images simultaneously while imaging. Beyond scale, another central advance of this work lies in the development and validation of a simple, end-to-end *in vivo* technological pipeline with physiological sensitivity to transcriptional activity and timestamp precision of 3.1–6.7 hours in median absolute error, compatible with expansion microscopy and RNA readouts. Since timestamps are delivered via chemical manipulation of drinking water, temporal uncertainty in timestamp induction may partially arise from individual variability in the timing and volume of voluntary water consumption. In principle, acute injection of timestamp-inducing substances could reduce this uncertainty by enabling precise control over delivery timing and dosage.

However, the injection procedure itself and the associated handling of the subject may risk evoking stress-related or somatosensory neural activity, and additional efforts are needed to characterize the extent to which such a procedure may influence the recorded cell activities and to develop protocols that minimize these confounds in data interpretation. Together, these new capabilities enabled high-throughput construction of a spatiotemporal, brain-wide atlas of memory under contextual fear conditioning and recall, a widely used paradigm in learning and memory research, and established a scalable resource and generalizable framework applicable across behavioral paradigms and disease models.

In this work, we characterized and validated GLOBE using the XRI monomer design. We found the XRI monomer design is sufficient for days-scale recording durations, robustly forming tapes that records transcriptional activity across diverse brain regions. The original XRI recording system has been demonstrated to record transcriptional activity along an absolute time axis in single cultured neurons, empowered by single-tape length-to-time calibration via the fractional cumulative expression of the structural monomer per tape^37^. Although the iCreERT/4-OHT system works robustly with XRI *in vivo*, due to its irreversible nature, it can provide only a single timestamp along the tape^37^(**Extended Data Fig. 4**), whereas the rtTA/Dox system used in this work can provide multiple timestamps due to its reversibility (**Extended Data Fig. 8)**. Future work could leverage rapidly evolving monomer designs for protein tape recorders and equip GLOBE recording with even more powerful capabilities, such as weeks-long recording duration and simultaneous recording of multiple kinds of cellular activities in the same cells^38^, beyond what we demonstrated here.

The *Fos* promoter has been widely used as an activity marker in studies of learning and memory in mammalian brains^19,24–28,30–35,56^. Nevertheless, promoter-based readout alone does not capture transcriptional regulatory events beyond promoter control or non-transcriptional mechanisms that shape *Fos* protein levels and mediate brain functions^49,57–61^. Due to the lack of these additional regulatory mechanisms, *Fos* promoter-based readout may exhibit slower signal decay kinetics after activation, with *Fos* promoter activity lasting even multiple days in certain neurons as observed in our study and in previous studies^14,25,47–49^, than the hours-scale dynamics of *Fos* protein abundance and the sub-hour-scale dynamics of *Fos* mRNA abundance usually observed in neurons *in vivo*^62^, limiting its use in parsing behavioral events to ones that are well separated in time. Transgenic and knock-in mouse lines with *Fos* reporters are powerful tools for labeling *Fos*-positive cells^17,18,25,32,63^. GLOBE recording, by contrast, employs AAV-delivered, temporally continuous recorders, offering complementary flexibility for studies in animal models without transgenic or knock-in *Fos* reporters and for modular targeting of selected cell populations through construct engineering or targeted AAV injection.

Although behavioral manipulations and control experiments provide evidence that the recorded single-cell activity dynamics are behaviorally relevant, this study remains observational at the cellular level. Establishing mechanistic causality will require direct perturbation of defined cellular ensembles to test whether these dynamics are necessary or sufficient for memory formation and retrieval^64,65^. Future work may combine GLOBE recording with tools for large-scale molecular or activity perturbation of cell ensembles to causally dissect how cellular activity transforms behavioral, pharmacological, or environmental inputs into behavioral and physiological outcomes. Such efforts may help establish the causal functional roles of distinct cell ensembles, cell types, neural circuits, and brain regions in brain function^5^.

The distinct functional roles of many gene regulation dynamics beyond *Fos*, such as immediate early genes *Arc*^66^, *Npas4*^67^, *Egr1* (also called *Zif268*)^68^, and transcriptional factors, such as CREB^69^, have been reported across mammalian brains. Future work could integrate additional transient and long-lasting regulatory activities into protein tape recorders for GLOBE recording of these processes, ideally simultaneously within the same cells^38^, to dissect their distinct roles across the brain. Future work may also combine GLOBE recording with *in situ* molecular, structural, lineage and connectomic analyses to correlate spatiotemporal signatures of cell activity with genetic, epigenetic, transcriptomic, proteomic, cell-lineage, anatomical, circuit-level, cellular, and subcellular features, among other relevant molecular and spatial properties, to enable more comprehensive multimodal analyses beyond those demonstrated in this work.

## 4 Methods

### Molecular cloning

The DNAs used in this work were mammalian-codon optimized, synthesized by Epoch Life Science, and then cloned into AAV plasmid backbones for viral packaging and *in vivo* delivery. For constructs used for AAV9 preparation, pAAV-UbC was used to drive the constitutive expression of the structural monomer XRI-HA. pAAV-Syn-ERT2-iCre-ERT2 and pAAV-UbC-FLEX were used for the Cre-dependent expression of XRI-FLAG^37^. pAAV-Syn and pAAV-TRE3G (TakaraBio) were used to express rtTA3G (TakaraBio) and Dox-based timestamp monomer XRI-FLAG, respectively, for Dox-dependent expression^38^. For activity-dependent signal monomer expression, XRI-V5 was placed under the control of the *Fos* promoter (pAAV-Fos). In the signal monomer construct, a protein-destabilizing PEST sequence was fused in-frame to the C-terminus of the coding sequence, and an mRNA-destabilizing element was inserted downstream of the stop codon, resulting in XRI-V5-P1N4, as done previously^70,71^. For constructs used for AAV-PHP.eB and AAV.CAP-B10 preparation, XRI-HA was driven by the ubiquitous CMV-β-actin-intron-α-globin hybrid promoter (pAAV-CAG), with the WPRE sequence inserted in the 3’UTR to enhance the expression level.

### Animals and mouse surgery

All procedures in this work involving animals were conducted in accordance with the United States National Institutes of Health Guide for the Care and Use of Laboratory Animals, and were reviewed and approved by the University of Michigan Institutional Animal Care & Use Committee. Mice were maintained on a 12-h light-dark cycle (lights on at 06:00 EST) in a temperature-controlled environment at 22 ± 1 °C, with a relative humidity of 30–50%. Mice were individually housed post-surgery and throughout the remainder of the experiment. Experiments were conducted using male mice. For behavioral procedures, the number of mice per experiment was determined based on expected variance and effect size from previous studies^24,67^; no statistical method was used to predetermine sample size; mice were randomly allocated to either the experimental or control groups; experimenters were not blinded to group identity.

### Stereotactic injection

C57BL/6J mice (2-5 months of age; male; Jackson Laboratory) were anesthetized with 5% isoflurane during induction and placed on a heating pad in a stereotactic frame (RWD Instruments) with 1.5-2% isoflurane throughout surgery to maintain deep anesthesia. Ophthalmic ointment was applied to both eyes. Hair was removed with a hair removal cream, and the surgical site was cleaned with ethanol and betadine. Following this, an incision was made to expose the skull. Craniotomy was performed by drilling through the skull above the injection sites using a 0.5 mm diameter drill bit. The AAV mixture was injected using a pulled glass capillary with a pressure microinjector (RWD) at a rate of 100 nL/min. Following injection, the needle remained at each target site for 5 min to facilitate AAV diffusion into brain tissue. Each site received 350 nL of AAVs, unless specially noted. After surgery, mice received 5 mg/kg carprofen i.p. and were placed on a heating pad for recovery.

AAV mixture was prepared by mixing AAV stocks (serotype AAV9; UNC NeuroTools) at defined ratios to control the relative expression levels of each construct. The final viral genome copy (GC) of the structural monomer (XRI-HA) AAV in the mixture is 10-fold and 100-fold greater than those for the timestamp (XRI-FLAG) AAV and the signal monomer (XRI-V5) AAV, respectively, to ensure that the tapes are predominantly composed of structural monomers. A mixture of 1.01 μL included: AAV9-UbC-XRI-HA (stock titer, 1.58 × 10^13^ GC/mL; final volume of 0.4 μL and final viral genome copies of 6.32 × 10^9^ GC), AAV9-Fos-XRI-V5-P1N4 (stock titer, 3.14 × 10^13^ GC/mL; diluted with DPBS at 20 times, final volume of 0.04 μL and final viral genome copies of 6.32 × 10^7^ GC), AAV9-hSyn-rtTA3G (stock titer, 3.56 × 10^12^ GC/mL; final volume of 0.44 μL and final viral genome copies of 1.58 × 10^9^ GC), AAV9-TRE3G-XRI-FLAG (stock titer, 4.87 × 10^12^ GC/mL; final volume of 0.13 μL and final viral genome copies of 6.32 × 10^8^ GC).

Stereotactic injection coordinates were defined relative to bregma (AP, ML) and brain surface (DV).

For brain-wide AAV delivery, the total volumes of the AAV mixture injected into each site are: (1) AP 1.7 mm; ML ±0.35 mm; DV, -1.6 mm; 0.3 μL at each site; (2) AP 0.98 mm; ML ±0.35 mm; DV -1.6 mm; 0.3 μL at each site; (3) AP -1.35 mm; ML ±0.3 mm; DV -0.3, -1.5, -3, -4.5 mm; 0.45 μL at each site; (4) AP -1.4 mm; ML ±3.3 mm; DV -1.5, -4 mm; 0.4 μL at each site; (5) AP -2 mm; ML ±1.5 mm; DV -1.5, -3.5 mm; 0.5 μL at each site.

For hippocampal AAV delivery, AAV mixture was injected at AP -2 mm; ML +/- 1.5 mm; DV -1.5 mm; with a final volume of 0.8 μL.

For **Fig. 2h-l**, a final volume of 0.45 μL AAV mixture was injected to the hippocampus at AP -2 mm; ML +/- 1.5 mm; DV -1.5 mm: AAV9-UbC-XRI-HA (stock titer, 1.58 × 10^13^ GC/mL; final volume of 0.4 μL and final viral genome copies of 6.32 × 10^9^ GC), AAV9-Fos-XRI-V5-P1N4 (stock titer, 3.14 × 10^13^ GC/mL; diluted with DPBS at 20 times, final volume of 0.04 μL and final viral genome copies of 6.32 × 10^7^ GC), AAV9-Fos-GFP (stock titer, 4.70 × 10^13^ GC/mL; final volume of 0.13 μL and final viral genome copies of 6.32 × 10^9^ GC).

For experiments in **Extended Data Fig. 4**, the AAV mixture for injection was prepared by mixing the AAV stocks (serotype AAV9; UNC NeuroTools) as the following final viral genome copies: AAV9-UbC-XRI-HA (stock titer, 1.58 × 10^13^ GC/mL; final viral genome copies of 6.32 × 10^9^ GC), AAV9-Syn-ERT2-iCre-ERT2 (stock titer, 5.51 × 10^13^ GC/mL; final viral genome copies of 6.32 × 10^9^ GC), AAV9-UbC-FLEX-XRI-FLAG (stock titer, 3.68 × 10^13^ GC/mL; final viral genome copies of 6.32 × 10^9^ GC), and injected to the following sites: (1) AP -2 mm; ML ±1.5 mm; DV, -1.5 mm; (2) AP -1.4 mm; ML ±3.3 mm; DV, -4.2 mm; (3) AP +1.7 mm; ML ±0.35 mm; DV, -1.6 mm; (4) AP -1.5 mm; ML ±0.35 mm; DV, -3 mm; (5) AP -1.25 mm; ML ±0.3 mm; DV, -5.2 mm; (6) AP -1.55 mm; ML ±3.2 mm; DV, -4.2 mm.

### Intravenous injection

C57BL/6J mice (2-5 months of age; male; Jackson Laboratory) were anesthetized with 5% isoflurane during injection. Intravenous delivery of AAV-PHP.eB (stock titer, 6.85 × 10^12^ GC/mL; diluted with DPBS, final viral genome copies, 3.43 x 10^11^ GC) or AAV.CAP-10 (stock titer, 3.30 × 10^12^ GC/mL; diluted with DPBS, final viral genome copies, 1.65 x 10^11^ GC) for the expression of the structural monomer (XRI-HA) was performed via retro-orbital injection. Following injection, ophthalmic ointment was applied to the injected eye.

### Doxycycline oral delivery in mice

Doxycycline hyclate (Sigma) was dissolved in standard drinking water at 1 mg/mL, supplemented with 5% (w/v) sucrose. Standard drinking water in home cages was replaced with this doxycycline water. Following the treatment period, doxycycline water was replaced with standard drinking water to terminate doxycycline delivery.

### 4-OHT injection in mice

4-OHT (Sigma) was dissolved in 100% ethanol (Sigma) at 100 mg ml^−1^ by vortexing for 5 min. Next, the solution was mixed with corn oil (Sigma) to obtain a final concentration of 10 mg ml^−1^ 4-OHT by vortexing for 5 min and then sonicating for 30–60 min until the solution was clear. The 10 mg ml^−1^ 4-OHT solution was then loaded into syringes and administered to mice via i.p. injection at 40 mg kg^−1^.

### Behavioral procedures

Behavioral procedures were performed during the light phase of the light-dark cycle. Mice were transported via wheeled cart to and from the behavior room individually in covered cages. Floors of chambers were cleaned before and between animal experiments. Freezing behavior was assessed using the VFreeze software (Med Associates). Contextual fear conditioning (CFC) and recall experiments were performed in the context of a 64 x 36 x 79 cm camera-monitored fear conditioning chamber (Med Associates, NIR VFC System NIR-022MD) with grid floors, neutral white lighting, and scented with 1% acetic acid. For CFC, mice were allowed to explore the context in the conditioning chamber for a 180 s baseline period, followed by three presentations of a foot shock (0.8 mA, 1 s duration) delivered at a 60 s inter-shock interval (shock onsets at 180 s, 240 s, and 300 s). Following the final shock, mice remained in the chamber for a 120 s post-shock period. Mice returned to their original home cages and housing room following the procedure. For recall, mice were allowed to explore the same context as in CFC for 420 s, with no foot shock delivered. Mice returned to their original home cages and housing room following the procedure.

### Histology

Mice were perfused transracially with 1× PBS followed by 4% paraformaldehyde (Electron Microscopy Sciences) in 1× PBS. The brain was gently extracted from the skull and postfixed in 4% paraformaldehyde in 1× PBS overnight at 4 °C. The brain was then stored in 1× PBS at 4 °C until slicing. The brain was sectioned coronally at 30-µm using a vibratome (Leica VT1000 S) and then stored in 1× PBS at 4 °C until immunofluorescence staining.

### Immunofluorescence of brain tissue

Brain tissue sections were blocked overnight at 4 °C in MAXBlock blocking medium (Active Motif) supplemented with 0.5% Triton X-100 (Thermofisher Scientific) and 100 mM Glycine, followed by four washes for 15 min each at RT in MAXWash washing medium (Active Motif). Tissues were then incubated with primary antibodies diluted in MAXbind staining medium (Active Motif) supplemented with 0.5% Triton X-100 shaking overnight at 4 °C, and then washed in MAXWash washing medium three times for 15 min each at RT. Next, tissues were incubated overnight at 4 °C, protected from light, with fluorescent secondary antibodies and NeuroTrace Blue Fluorescent Nissl Stain (Invitrogen), both diluted in MAXbind staining medium supplemented with 0.5% Triton X-100.Tissues were then washed three times in MAXWash washing medium for 15 min each at room temperature and stored in 1× PBS at 4 °C until imaging.

### Antibodies and stains

Primary antibodies (1:500): anti-HA (Cell Signaling Technology 3724), anti-V5 (Invitrogen R960-25), anti-FLAG (Sigma F1804). Fluorescent secondary antibodies (1:500): Goat anti-Rabbit IgG(H+L) Alexa Fluor Plus 488 (Invitrogen A32731), Goat anti-Mouse IgG2a Alexa Fluor 546 (Invitrogen A21133), and Goat anti-Mouse IgG1 Alexa Fluor 647 (Invitrogen A21240). Nissl stain: NeuroTrace 435/455 Blue Fluorescent Nissl Stain (1:1000, Invitrogen N21479).

Additional details of primary antibodies, secondary antibodies, stains and other reagents used in this study are listed in **Supplementary Table 1**.

### Fluorescence microscopy

Fluorescence microscopy was performed using a spinning disk confocal microscope (Yokogawa CSU-W1 Confocal Scanner Unit on a Nikon Eclipse Ti2 inverted microscope) equipped with a 40X water immersion objective (NA 1.15; Nikon MRD77410), a 4X objective, a 10X objective, a Hamamatsu ORCA-Fusion BT sCMOS camera controlled by the NIS-Elements AR software, and laser/filter sets for 405 nm, 488 nm, 561 nm and 640 nm optical channels. Automated multi-channel, multi-location volumetric imaging was conducted at a 0.4µm per Z-step under the 40X objective. Imaging parameters remained consistent across all samples within each experimental set. The brain-wide readout from sectioned brain tissues took up to 177 hours of imaging per mouse brain, corresponding to an average imaging readout speed of approximately 2.9 seconds per cell.

### Brain atlas registration and associated imaging

For the brain-wide dataset, individual FOVs were manually aligned to the Allen Common Coordinate Framework version 3 (CCFv3) atlas^46^ based on anatomical landmarks, including nuclear density, ventricles, and tissue boundaries. After registration, each FOV was assigned to its corresponding anatomical region.

### Expansion microscopy of brain tissue slices

After fixation, the brain was sectioned coronally at 50-µm using a vibratome (Leica VT1000 S). Tissues were then incubated overnight at 4 °C with gentle shaking in permeabilization buffer containing 0.5 mM acrylic acid N-hydroxysuccinimide easter (AAX, Sigma A8060) and 10% (w/v) CHAPS (Sigma C3023) in MES (Sigma M3671) buffered saline at pH 6. Tissues were then transferred to pH 8.3 HEPES-buffered saline (Thermofisher Scientific J61239.AP) for neutralization and incubated at 4 °C for 6 h to activate AAX crosslinking. The monomer solution used to cast the hydrogel consisted of 5.3% (w/v) sodium acrylate (SA; Sigma 408220), 4% (w/v) acrylamide (AA; Bio-Rad 1610140), 0.02% (w/v) N,N′-methylenebisacrylamide (Bis; Bio-Rad 1610142), and 2% (v/v) N,N-dimethylacrylamide (DMAA; Sigma 274135) in 1× PBS. The solution was degassed under vacuum in 7 mL glass vials prior to use and kept on ice in between degassing ultrasound bursts. VA-044 (FUJIFILM, LB-VA044) polymerization initiator was added to a final concentration of 0.5% (w/v), and tissues were incubated in the monomer solution overnight at 4°C with gentle shaking. Tissues were then mounted onto glass slides with 50 µm spacers, overlaid with gelling solution, and sealed with a coverslip. Polymerization was carried out at 37 °C for 6 h in a humidified chamber. Gel-embedded tissues were excised and transferred to new glass vials. Clearing was performed by incubating tissues in softening buffer containing 20% (w/v) SDS (Sigma 436143), 100 mM β-mercaptoethanol (Millipore 444203), 25 mM EDTA (Invitrogen), 0.5% Triton X-100 (Sigma), and 50 mM Tris (Invitrogen, pH 8) at 95 °C for 2 h using a thermocycler. Samples were subsequently washed in washing buffer, which contained 300 mM betaine (Sigma B2754), 4% bile (Millipore B8756), and 2% β-cyclodextrin (Sigma C4767) in HEPES buffer pH 7.8, three times (20 min each) at 37 °C with gentle shaking. Tissues were incubated with primary and secondary antibodies (both at 1:100 dilution) as described above. For the re-embedding step, the same gel formulation was used with 150 µm spacers. After the second gelation, tissues were subjected to a second round of immunostaining and imaged.

### RCA-based RNA FISH

Mice were perfused transcardially with RNAse-free 1× PBS followed by 4% paraformaldehyde (Electron Microscopy Sciences) in 1× PBS, and all subsequent tissue processing steps were performed under RNase-free conditions.

Sectioned brain tissues were washed three times in PBS at room temperature and incubated in PBST containing 0.25% Tween-20 in PBS for 20 min at room temperature. Sections were permeabilized with 0.1% Triton X-100 in 1× PBS for 1 h at room temperature, followed by three washes in PBST for 20 min each. Tissues were then incubated with 0.5 mM AAX in 1× PBS overnight at 4 °C. After incubation, sections were washed three times in 1× PBS for 15 min each. For gelation, tissues were incubated overnight at 4 °C in monomer solution containing 4% (w/w) acrylamide, 0.2% (w/w) N,N′-methylenebisacrylamide and 0.1% (w/v) VA-044 in 1× PBS. Tissues were then placed within 50-µm spacers and polymerized at 37 °C for 3 h. After gelation, samples were washed three times in 1× PBS for 10 min each. Proteinase K digestion was performed at 37 °C for 30 min, after which sections were washed three times in PBS for 20 min each at room temperature. Sections were blocked in the hybridization buffer for 30 min at room temperature. Padlock probes and amplification primers were diluted in hybridization buffer to final concentrations of 20 nM and 200 nM, respectively, and applied to the sections overnight at 37 °C. The hybridization buffer contained 0.1 mg/mL salmon sperm DNA, 50 mM KCl, 20% formamide, 20 µg/mL BSA, 20 µg/mL yeast tRNA and 1× Ampligase buffer. The buffer was mixed thoroughly and kept on ice or stored at −20 °C until use. After hybridization, sections were washed four times in a hybridization washing buffer containing 10% formamide in 2× SSC at 37 °C for 20 min each, followed by two washes in PBST at room temperature for 30 min each. SplintR ligation was performed for 3 h at room temperature in a 300-µL reaction containing 6 µL SplintR ligase (5 U/µL; final concentration, 0.1 U/µL; NEB M0375S), 30 µL 10× ligase buffer, 3 µL BSA (20 mg/mL; final concentration, 0.2 mg/mL; NEB B9200S), 1 µL RNase inhibitor (Sigma 3335399001) and nuclease-free water to a final volume of 300 µL. Sections were then washed three times in PBST containing 0.1% Tween-20 in 1× PBS for 20 min each at room temperature. Rolling circle amplification was performed overnight at 30 °C in a 100-µL reaction containing 2 µL Phi29 DNA polymerase (10 U/µL; final concentration, 0.2 U/µL; NEB M0269S), 10 µL 10× reaction buffer, 1 µL BSA (20 mg/mL; final concentration, 0.2 mg/mL), 1 µL RNase inhibitor, 2.5 µL dNTPs (10 mM each; final concentration, 0.25 mM; Invitrogen 18427088) and nuclease-free water to a final volume of 100 µL. After amplification, sections were washed three times in PBST at room temperature for 20 min each. For imaging, sections were incubated with fluorophore-conjugated imager probes and imaged by confocal microscopy. For sequential imaging cycles, fluorophore-conjugated probes were stripped with 60% formamide in 1× PBS for 10 min at room temperature, followed by three washes in 1× PBS for 15 min each. Subsequent cycles of fluorophore-conjugated probe hybridization and imaging were performed as described above.

### Software for image analysis

Image analysis was performed in Python, ImageJ (National Institutes of Health), and napari (napari contributors; doi:10.5281/zenodo.3555620).

### Computational readout of protein tape recordings and cell boundaries from microscopy images of mouse brain tissue

We developed GLOBE Tape Reader, a scalable computational pipeline for brain-wide readout of protein tapes and cell morphology from multi-channel volumetric confocal microscopy images of mouse brain tissue. GLOBE Tape Reader builds on our previously published Tape Reader framework^38^, which established the core workflow for tape segmentation, skeletonization, signal extraction, and bilateral readout. However, the original Tape Reader was optimized for hippocampus datasets and did not generalize sufficiently to the brain-wide dataset from GLOBE recording. We therefore redesigned the pipeline for brain-wide analysis by expanding the training data across diverse brain regions, upgrading the segmentation backbone from U-Net to MedNeXt-S, optimizing GPU-based parallel inference and batched signal extraction, and incorporating morphology-, soma-, signal-, and symmetry-based quality control criteria tailored to the brain-wide dataset.

GLOBE Tape Reader takes as input multi-tile ND2 acquisitions containing four-channel volumetric images: 405 nm Nissl stain, 488 nm structural monomer, 561 nm signal monomer, and 647 nm timestamp monomer. Stage coordinates and voxel dimensions were read from ND2 metadata for physical-coordinate analysis. The workflow consisted of four main steps. First, protein tapes were segmented from the 488 nm channel using a MedNeXt-S 3D neural network implemented in PyTorch Connectomics^72,73^. The model predicted foreground, contour, and skeleton-aware distance-transform maps, which were decoded into tape instances using marker-based watershed. The model was trained on 12 manually annotated 3D volumes from diverse brain regions, with watershed parameters optimized on a held-out validation volume. Inference used sliding-window prediction, test-time augmentation, intensity clipping, and GPU-parallel tile processing. Second, predicted tape instances were converted into physical-coordinate point clouds, skeletonized by PCA-guided spline fitting, and sampled into standardized 1,000-point centerlines. Third, intensity profiles from all four channels were extracted along each skeleton using batched trilinear interpolation. The tape midpoint was optimized using the 647 nm timestamp channel to maximize bilateral symmetry, after which each tape was split into two 500-point side-arm profiles and scored by left–right Pearson correlation. Fourth, neuronal soma boundaries were segmented from the 405 nm Nissl channel using micro-SAM^74^, and tapes were assigned to parent cells by sampling soma labels along the skeleton. Final outputs included tape IDs, parent cell IDs, lengths, centroids, midpoint positions, symmetry scores, and four-channel intensity profiles. Tapes were retained for downstream analysis only if they satisfied soma-assignment, morphology, signal-detection, and bilateral timestamp-symmetry criteria.

The quality control criteria were applied as follows. Soma-assignment criteria required segmented tapes located inside neuronal soma, defined as having at least 10% of skeleton points with Nissl intensity greater than 100. Morphology criteria required tapes to have a geodesic length of ≤ 25 µm, a volume greater than 2 µm^3^, a z-span of ≤ 10 optical sections, and a skeleton PCA linearity score greater than 0.8. The PCA linearity score was defined as the fraction of variance in skeleton coordinates explained by the first principal component; values greater than 0.8 were considered indicative of tape-like morphology. The signal-detection criteria required tapes to exhibit detectable signal in structural monomer, Dox-dependent timestamp monomer, and signal monomer channels, with maximum intensities greater than 200 A.U., 200 A.U., and 120 A.U., respectively. The bilateral timestamp-symmetry criteria required Dox-dependent signals to be detected on both sides of the tape center with strong bidirectional symmetry, defined as both side-arm signals exceeding the central baseline and a left-right Pearson correlation coefficient greater than 0.9.

### Computational resources

Computation for training and applying GLOBE Tape Reader was performed on a Linux high-performance computing cluster. The tape segmentation model was trained using 4 NVIDIA A100 GPUs (40 GB memory each) with a batch size of 4 and an input patch size of 32 x 96 x 96 voxels. Training ran for 800 epochs, lasting roughly 5 hours. The end-to-end analysis pipeline was orchestrated as a four-stage SLURM job array with inter-stage dependencies: (1) tile extraction (CPU), (2) tape segmentation inference (GPU, ∼4 minutes per tile), (3) cell segmentation (GPU, ∼6 minutes per tile), and (4) tape analysis including skeletonization, signal extraction, and normalization (CPU with parallelization across tapes using joblib, ∼6-10 minutes per tile depending on tape density). Stages 2-4 were parallelized across tiles, enabling all tiles within an ND2 file to be processed simultaneously. Tape and cell segmentation inference each required one NVIDIA A100 GPU per tile. Per mouse brain, we processed up to 417 ND2 acquisitions comprising 3,038 individual imaging tiles from all regions of the brain, yielding segmented and analyzed protein tapes (7.5 TB of image data), within 878 hours of compute time, consisting of 488 gpu hours and 390 cpu hours, at an average computational readout time of 1.01 seconds per cell.

### Time recovery with TRE/Dox-based timestamp

To convert recorded signals into temporally resolved representations, we leveraged the robust ON and OFF waveform features of the Dox-dependent (FLAG) signal as timestamps. For each tape in the 15-day experiment, the onset of the rise of the FLAG signal, the onset of the decay of the FLAG signal, and the end of the FLAG signal were used as timestamps for Day 10, Day 11, and Day 15, respectively. Space-to-time transformation was performed by curve fitting the location coordinates of the timestamps along tape and the corresponding real world timepoints to a piecewise linear function directly connecting the timestamp points.

### Classification of temporal modes

*Fos* signal time courses (referred to as “traces”) with an absolute peak-amplitude below 0.2 were first classified as the non-responsive mode. The remaining traces were further classified using a supervised random forest model trained on manually annotated traces from randomly selected hippocampal neurons. The classifier was iteratively refined through repeated manual review of prediction outputs. The resulting trained classification model was then applied uniformly to all traces except for those already in the non-responsive mode.

### Statistical analysis

All statistical analysis was performed using the built-in statistical analysis tools in Prism (GraphPad) or Python. The details of each statistical analysis can be found in the figure legends. All statistical tests were two-sided.

## Acknowledgments

We thank Sara Aton, Lu Jiang, Catherine Kaczorowski, Michael Z. Lin, Shannon Moore, Geoffrey Murphy, Jennifer Murray, Henry Paulson, Katherine Stangis, Jim Stanis, and Natalie Tronson for discussion and technical support. Y.Y. acknowledges the Rackham Graduate Student Research Grant. J.G. is supported by National Science Foundation (NSF) Graduate Research Fellowship Program (2241144).

D.C. is supported by National Institutes of Health (NIH) NIAMS (1UC2AR082197-01) and NIMH (1R01MH134803-01A1). D.W. is NSF CAREER Award (IIS-2239688). C.L. and D.J.C. are supported by the Chan Zuckerberg Initiative Collaborative Pairs Pilot Project Award. C.L. is supported by the NIH Director’s New Innovator Award (DP2MH140133), the Glenn Foundation for Medical Research and American Federation for Aging Research Grant Award for Junior Faculty, the Whitehall Foundation, and the Klingenstein Fellowship Award in Neuroscience.

## Declarations

### Data availability

Plasmids and the corresponding sequence maps of constructs reported in this paper will be available at Addgene upon completion of the deposition process. Sequences of the structural, timestamp, and signal monomers will also be available at GenBank upon completion of the deposition process. The full brain-wide single-cell activity recording dataset is available at Zenodo (DOI: 10.5281/zenodo.20241204).

### Code availability

Source code for GLOBE Tape Reader can be found at https://github.com/LinghuLab/GLOBETapeReader.

### Author contribution

D.S. and C.L. conceived the concept of GLOBE and made high-level plans for this project. D.S., Y.H., W.C.J., and C.L. developed the experimental pipeline. Y.Y., T.-H.Z., P.L., M.G., and D.W. developed the computational pipeline for readout of protein tape recordings. D.S., Y.H., W.C.J., Y.Y., Y.W., J.L., L.Z. performed experiments. T.-H.Z., Y.Y., D.S., Y.H., P.L., J.G., M.G., F.L., A.D., and D.W. performed computational analysis. Y.H., X.N., M.-C.C., J.-C.H., and D.C. performed additional expansion microscopy and RNA readout. D.S., Y.H., Y.Y, B.K., D.J.C., D.W., and C.L. interpreted the data. D.S. and C.L. wrote the manuscript with the input from all authors. C.L. supervised the project.

---

**Extended Data Fig. 1.**
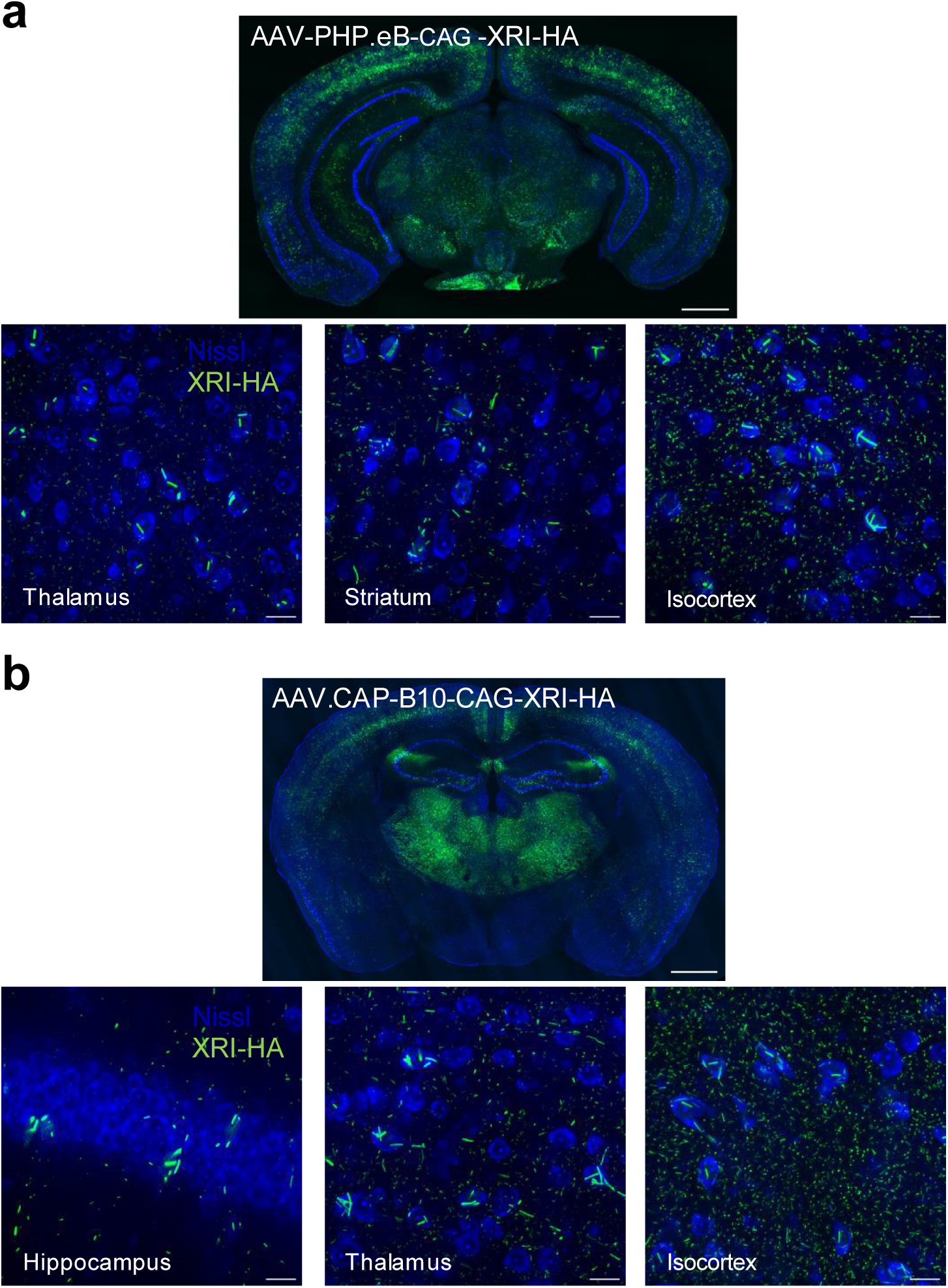
Brain wide expression of XRI via intravenous AAV delivery. **a**, Coronal section image (top) showing XRI expression following intravenous delivery of AAV-PHP.eB-CAG-XRI-HA, with representative confocal images (bottom) showing the morphology of XRI assemblies in selected brain regions. **b,** Coronal section image (top) showing XRI expression following intravenous delivery of AAV.CAP-B10-CAG-XRI-HA, and representative confocal images (bottom) showing the morphology of XRI assemblies in selected brain regions. Green, XRI via anti-HA immunostaining; blue, neuronal soma via Nissl staining. Scale bars, 20 μm.

**Extended Data Fig. 2.**
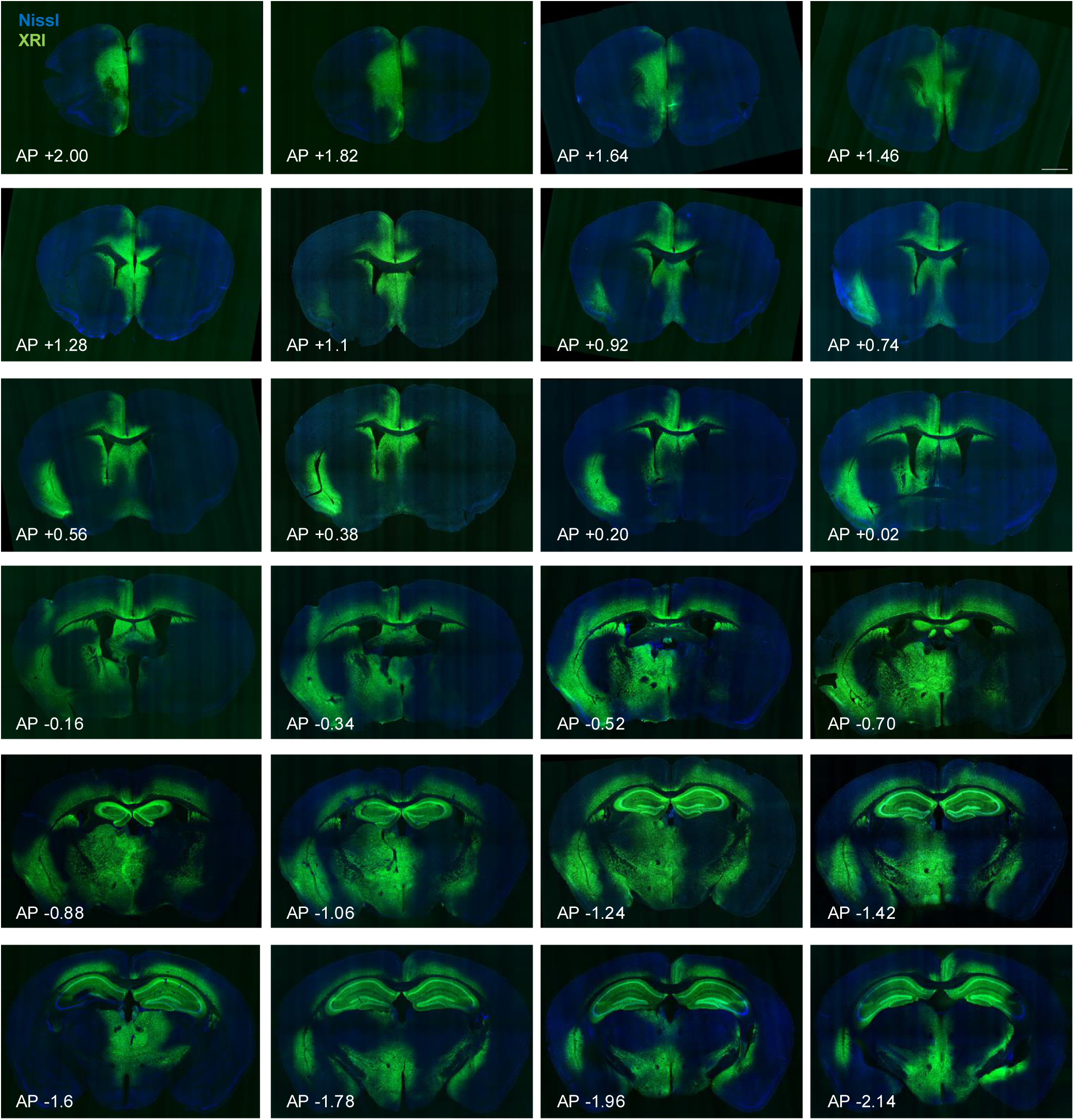
Brain-wide expression of XRI via intracranial AAV injection. Coronal sections images showing widespread XRI expression across the mouse brain, delivered by intracranial AAV injection of AAV9-UbC-XRI-HA. Sections are arranged from anterior to posterior (left-to-right, top-to-bottom) at 0.18 mm intervals. Anterior–posterior (AP) coordinates are indicated relative to bregma based on the Allen CCFv3 atlas. The brain was sectioned coronally at 30 µm thickness, with one section shown per six consecutive slices. Green, XRI via anti-HA immunostaining; blue, neuronal soma via Nissl staining. Scale bar, 1,000 μm.

**Extended Data Fig. 3.**
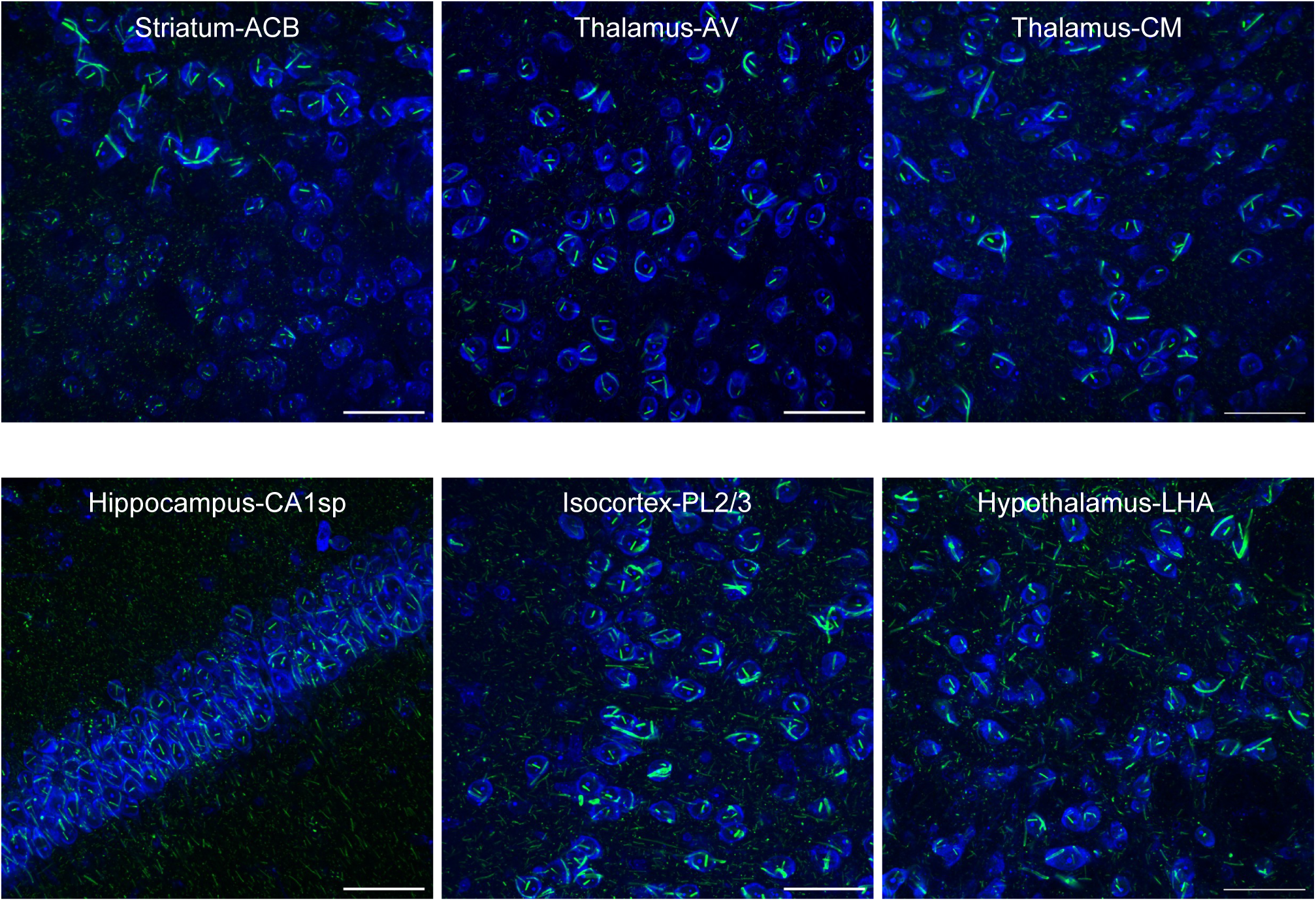
Expression and morphology of XRI assemblies across brain regions, via intracranial AAV injection. Representative confocal images showing the morphology of XRI assemblies across multiple brain regions, including striatum (ACB), thalamus (AV, CM), hippocampus (CA1sp), isocortex (PL2/3), and hypothalamus (LHA), delivered by intracrannial AAV injection of AAV9-UbC-XRI-HA. Green, XRI via anti-HA immunostaining; blue, neuronal soma via Nissl staining. Scale bars, 50 μm.

**Extended Data Fig. 4.**
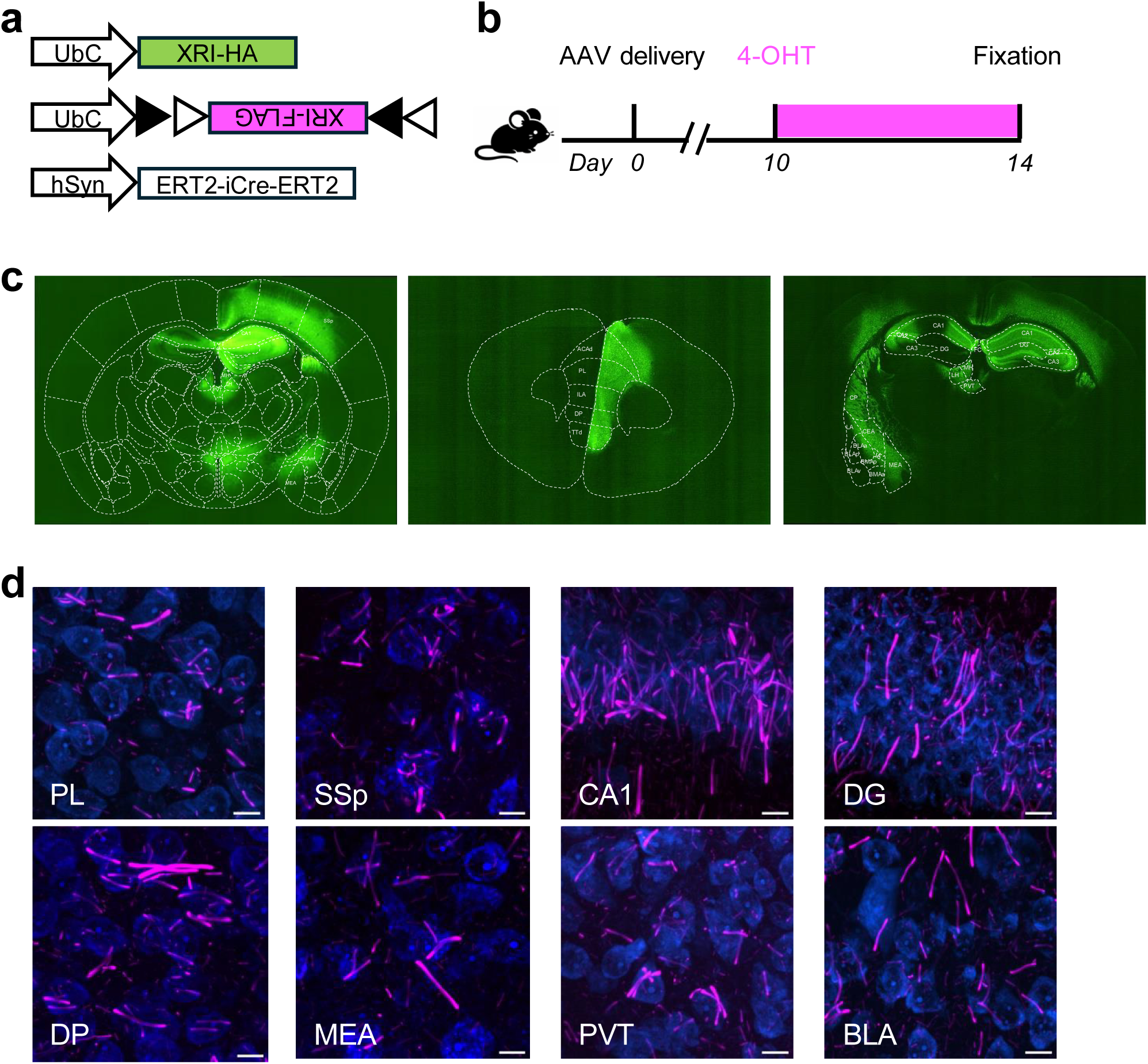
Validation of single cell temporal encoding across brain regions. **a, b**, Schematic of the AAV constructs (**a**) and experiment pipeline (**b**) for XRI structural monomer expression (UbC-XRI-HA), 4-OHT-inducible monomer expression (UbC-FLEX-XRI-FLAG), and Syn-ERT2-iCre-ERT2 expression. AAVs were injected to multiple regions of the mouse brain at day 0, followed by 4-OHT i.p. injection at day 10 and fixation at day 14. **c,** Representative coronal sections images showing XRI expression across the brain. **d,** Representative confocal images showing the bipolar distribution of 4-OHT-induced monomers along XRI tapes across brain regions, including prelimbic cortex (PL), primary somatosensory cortex (SSp), hippocampal CA1, dentate gyrus (DG), dorsal peduncular cortex (DP), medial amygdala (MEA), paraventricular thalamus (PVT), and basolateral amygdala (BLA). Scale bars, 10 μm.

**Extended Data Fig. 5.**
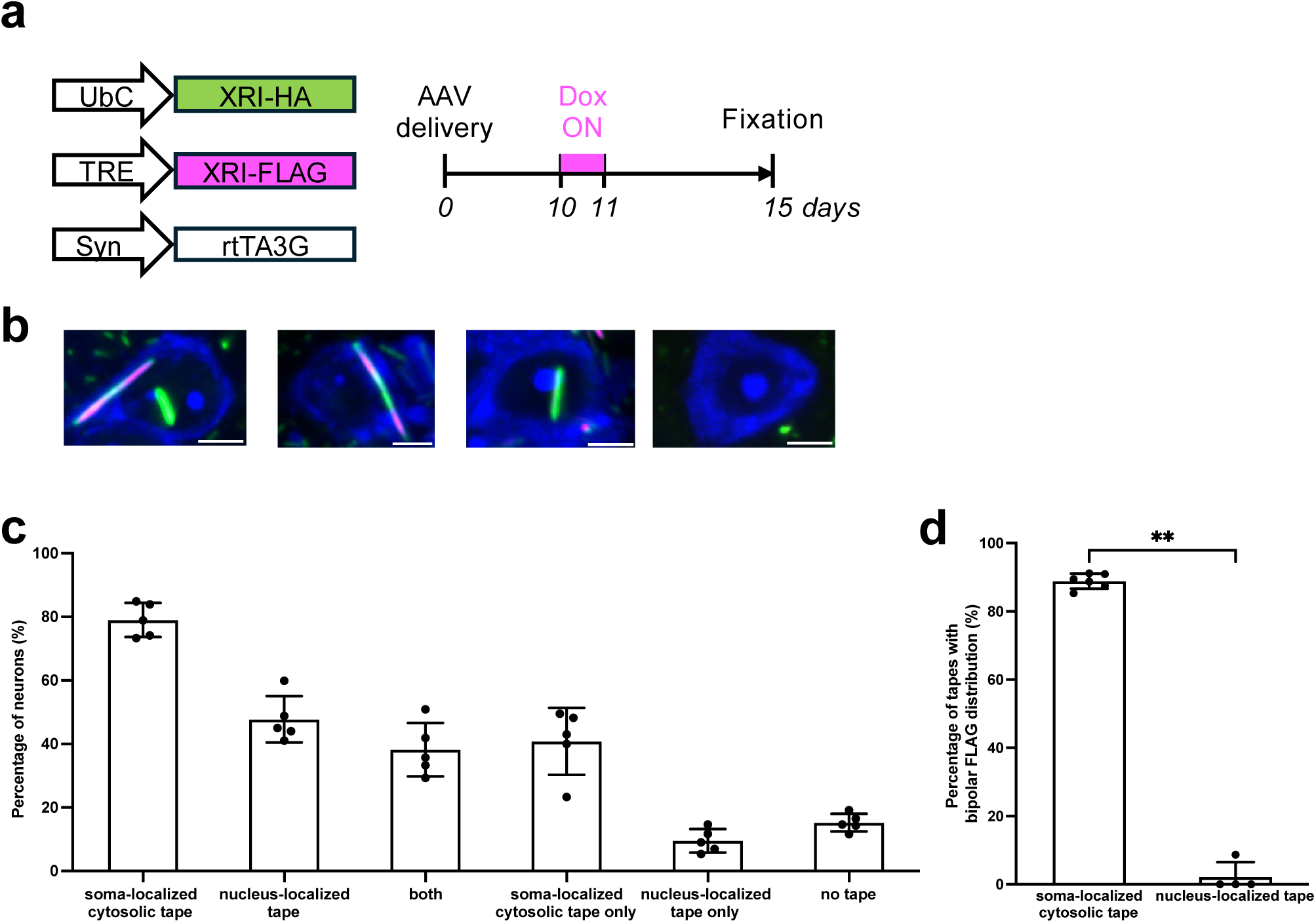
Subcellular localization of XRI tapes. **a**, Schematic of AAV constructs (top) and experimental pipeline (bottom) for XRI structural monomer expression (UbC-XRI-HA), Dox-dependent timestamp monomer expression (TRE-XRI-FLAG), and Syn-rtTA3G expression. AAVs were injected into mouse hippocampus at day 0, followed by Dox administration via drinking water from day 10 to day 11, and fixation at day 15. **b,** Confocal images of representative neurons with both soma-localized cytosolic tapes and nucleus-localized tapes, with soma-localized cytosolic tape only, with nucleus-localized tape only, and with no tapes. Green, structural monomer via anti-HA immunostaining; magenta, Dox-dependent timestamp monomer via anti-FLAG immunostaining; blue, neuronal soma via Nissl staining. Scale bars, 5 μm. **c**, Percentage of neurons with soma-localized cytosolic tapes, nucleus-localized tapes, both, soma-localized cytosolic tapes only, nucleus-localized tapes only, or no tapes. *n* = 56, 109, 120, 232, 86 neurons from 5 brain sections from 4 mice. **d,** Percentage of tapes with Dox-induced monomers localized at both ends of the tapes for soma-localized cytosolic tapes and for nucleus-localized tapes. **P = 0.01, two-sided Mann–Whitney U test.

**Extended Data Fig. 6.**
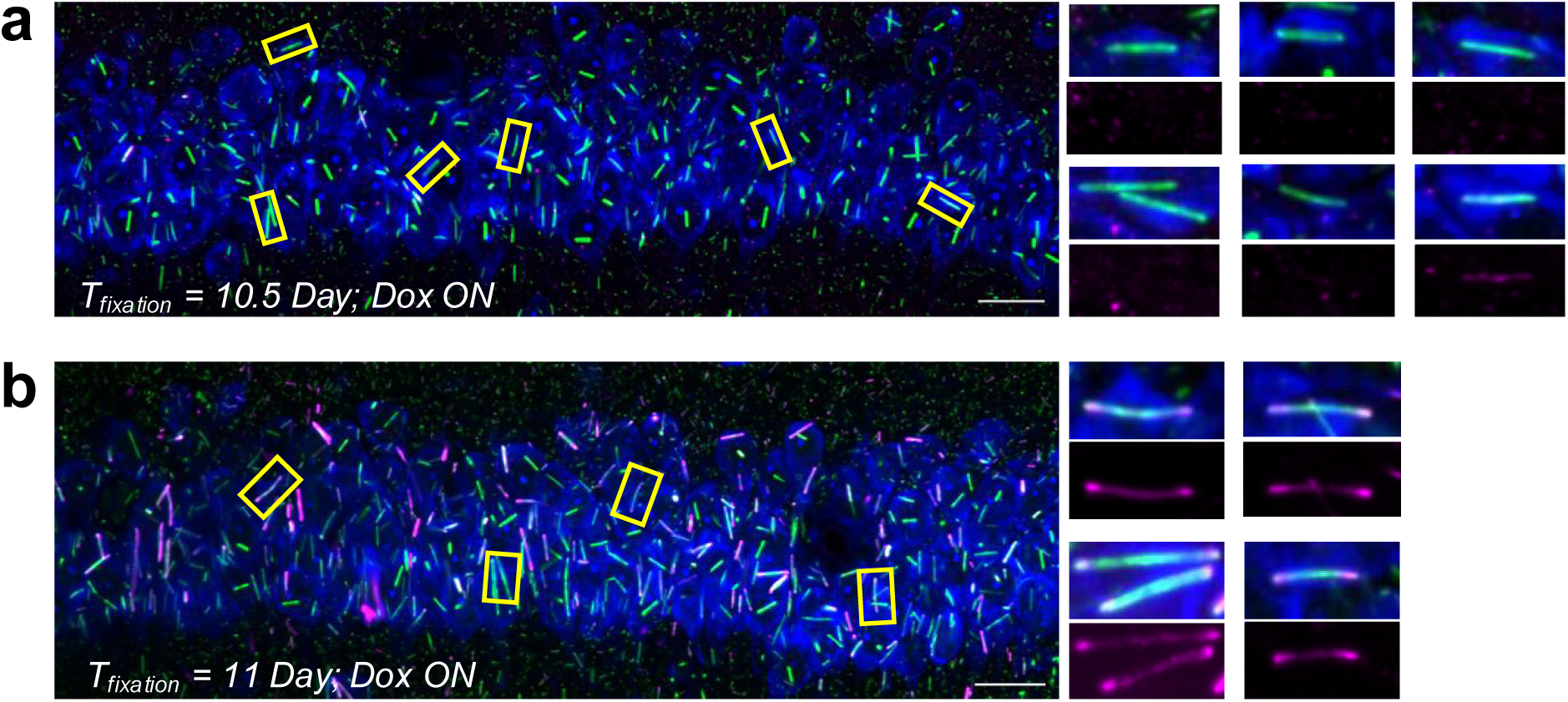
Confocal images of Dox-dependent signals following Dox administration. **a,b**, Representative confocal images of brain sections collected at day 10.5 (**a**) and day 11 (**b**). AAVs were injected into mouse hippocampus at day 0, followed by Dox administration via drinking water at day 10. Green, structural monomer via anti-HA immunostaining; magenta, Dox-dependent timestamp monomer via anti-FLAG immunostaining; blue, neuronal soma via Nissl staining. Yellow boxes indicate regions of interest, with corresponding enlarged views shown on the right. Scale bars, 20 μm.

**Extended Data Fig. 7.**
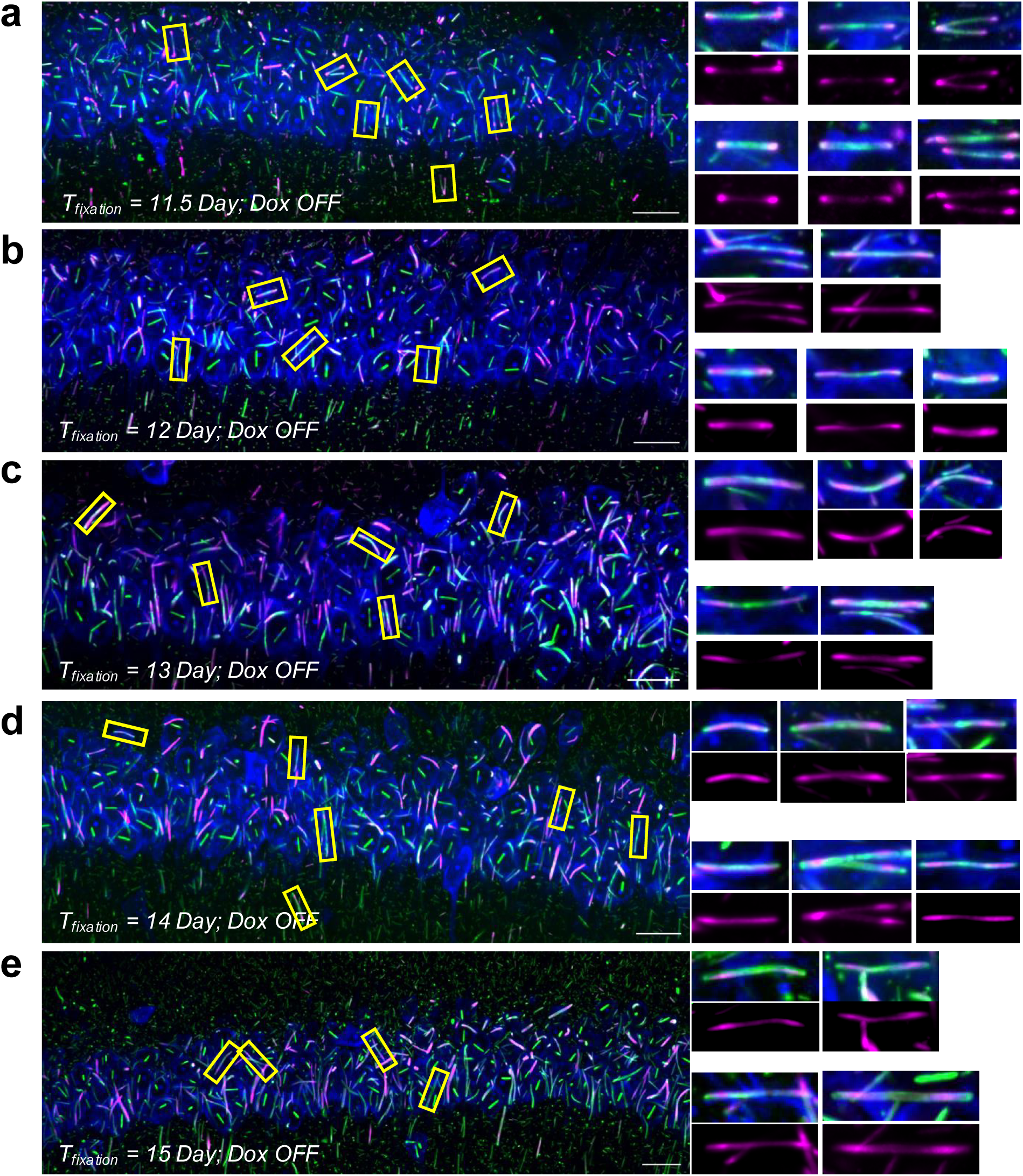
Confocal images of Dox-dependent signals following Dox withdrawal. **a–e**, Representative confocal images of brain sections collected at day 11.5 (**a**), day 12 (**b**), day 13 (**c**), day 14 (**d**), and day 15 (**e**). AAVs were injected into mouse hippocampus at day 0, followed by Dox administration via drinking water at day 10, and Dox withdrawal at day 11. Green, structural monomer via anti-HA immunostaining; magenta, Dox-dependent timestamp monomer via anti-FLAG immunostaining; blue, neuronal soma via Nissl staining. Yellow boxes indicate regions of interest, with corresponding enlarged views shown on the right. Scale bars, 20 μm.

**Extended Data Fig. 8.**
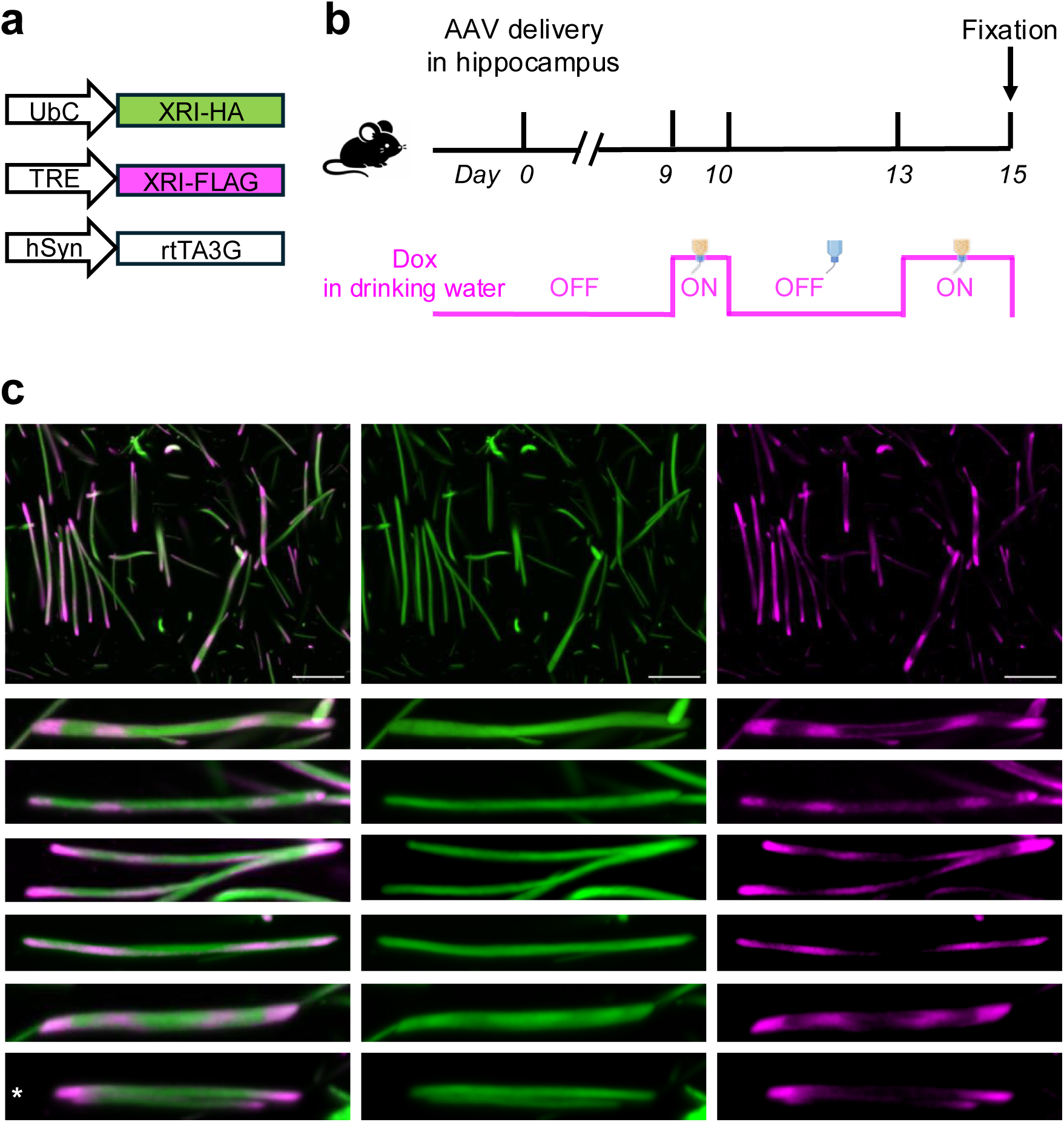
Timestamps induced by multiple Dox ON/OFF switching cycles visualized under expansion microscopy readout. **a,b**, Schematics of AAV constructs (**a**) and experimental design (**b**). AAVs were delivered into the hippocampus at day 0, followed by alternating Dox ON and OFF switches in drinking water at user-defined time periods and fixation at day 15. **c,** Expansion microscopy images showing structural monomer (green) and Dox-dependent timestamp monomer (magenta). Spatially distinct bands of timestamp monomers that correspond to multiple Dox ON and OFF switches are resolved along individual tapes. Bottom panels are enlarged views of representative tapes from this field of view. The asterisk marks a tape that failed to encode the full set of induced timestamps. Scale bars, 4.54 μm.

**Extended Data Fig. 9.**
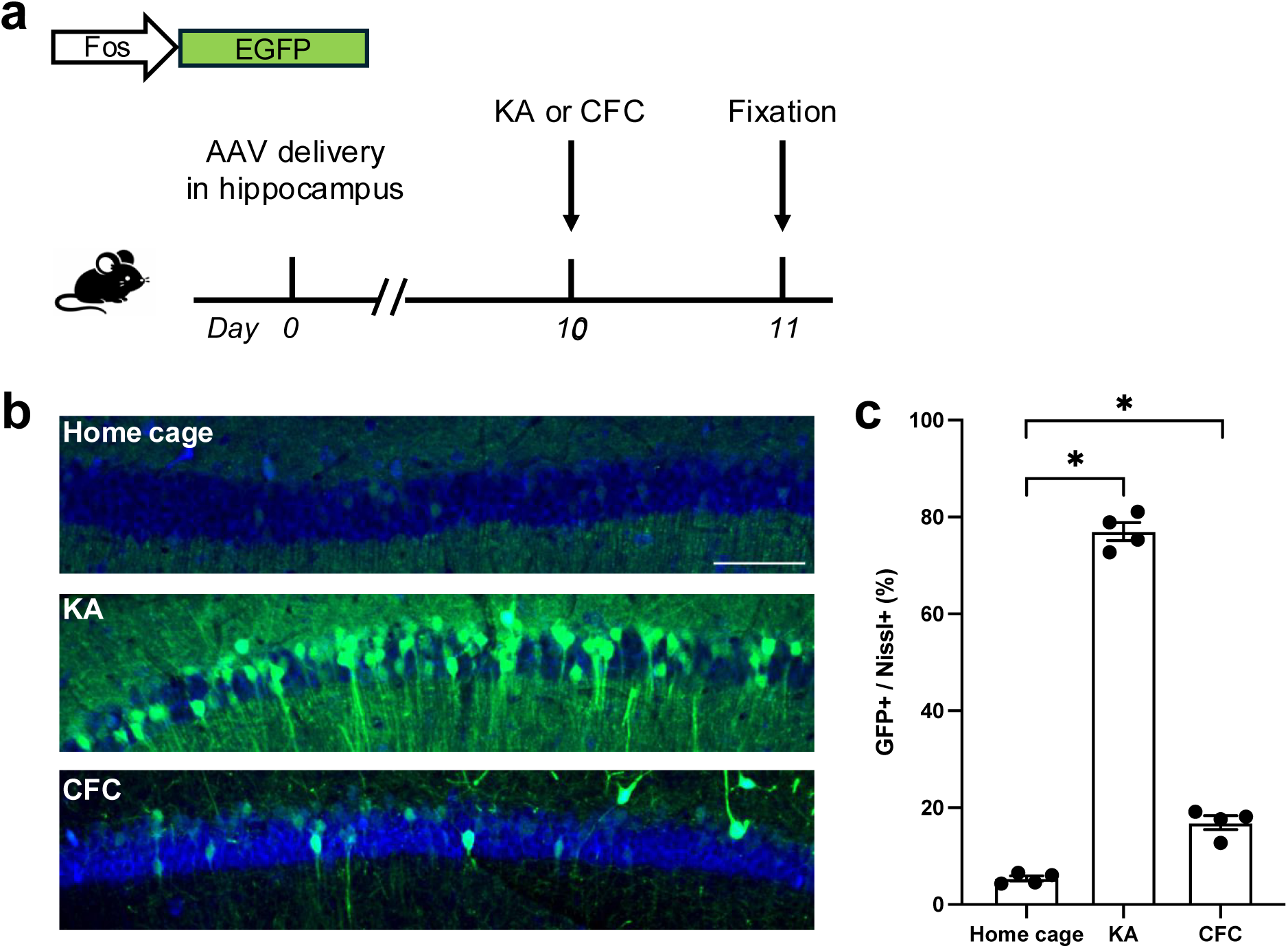
Validation of activity-dependent expression of *Fos*-GFP delivered by intracranial AAV injection. **a**, Schematic of AAV construct and experimental design. AAVs were delivered to the hippocampus at day 0, followed by contextual fear conditioning (CFC) or kainic acid (KA) i.p. injection at day 10 and fixation at day 11. **b,** Representative confocal images of hippocampal sections from mice under home cage, KA, and CFC conditions. Scale bars, 100 μm. **c,** Percentage of GFP-positive cells among Nissl-positive cells across conditions (home cage, KA, and CFC). Each point represents one brain section. *n* = 4 brain sections from 2 mice for each condition. *P < 0.05, two-sided Mann–Whitney U test.

**Extended Data Fig. 10.**
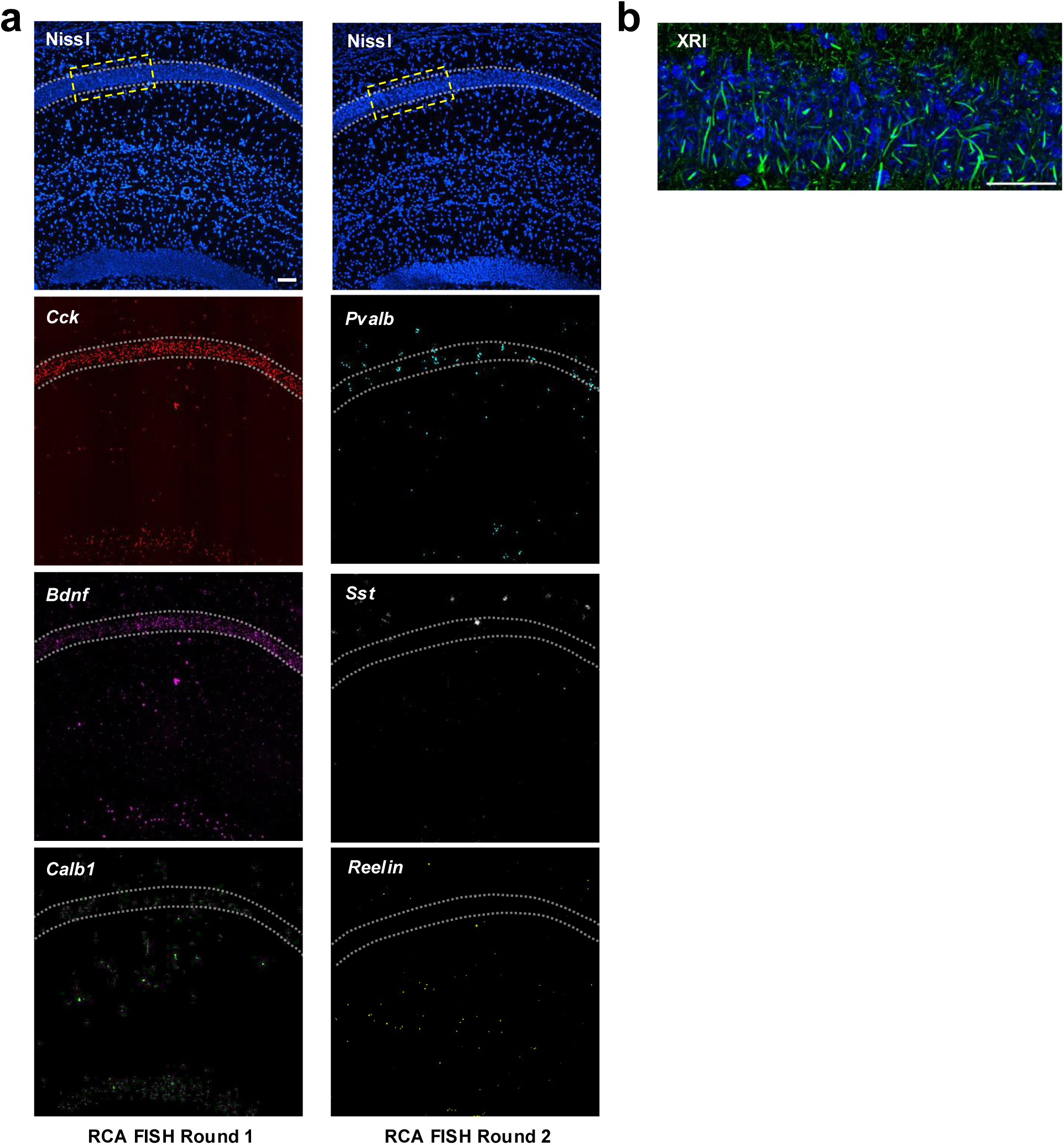
GLOBE is compatible with multiplexed, spatially-resolved RNA readout. **a**, Representative confocal images showing two sequential rounds of RNA readout in hippocampal tissue via multiplexed RCA-FISH. Dotted lines outline the CA1 pyramidal cell layer (stratum pyramidale). **b,** Representative confocal image showing XRI tapes in the region indicated by the dashed yellow box in **a**. Immunostaining of the structural monomer via the anti-HA antibody was performed after two rounds of RCA-FISH. Scale bars, 62.5 μm.

**Extended Data Fig. 11.**
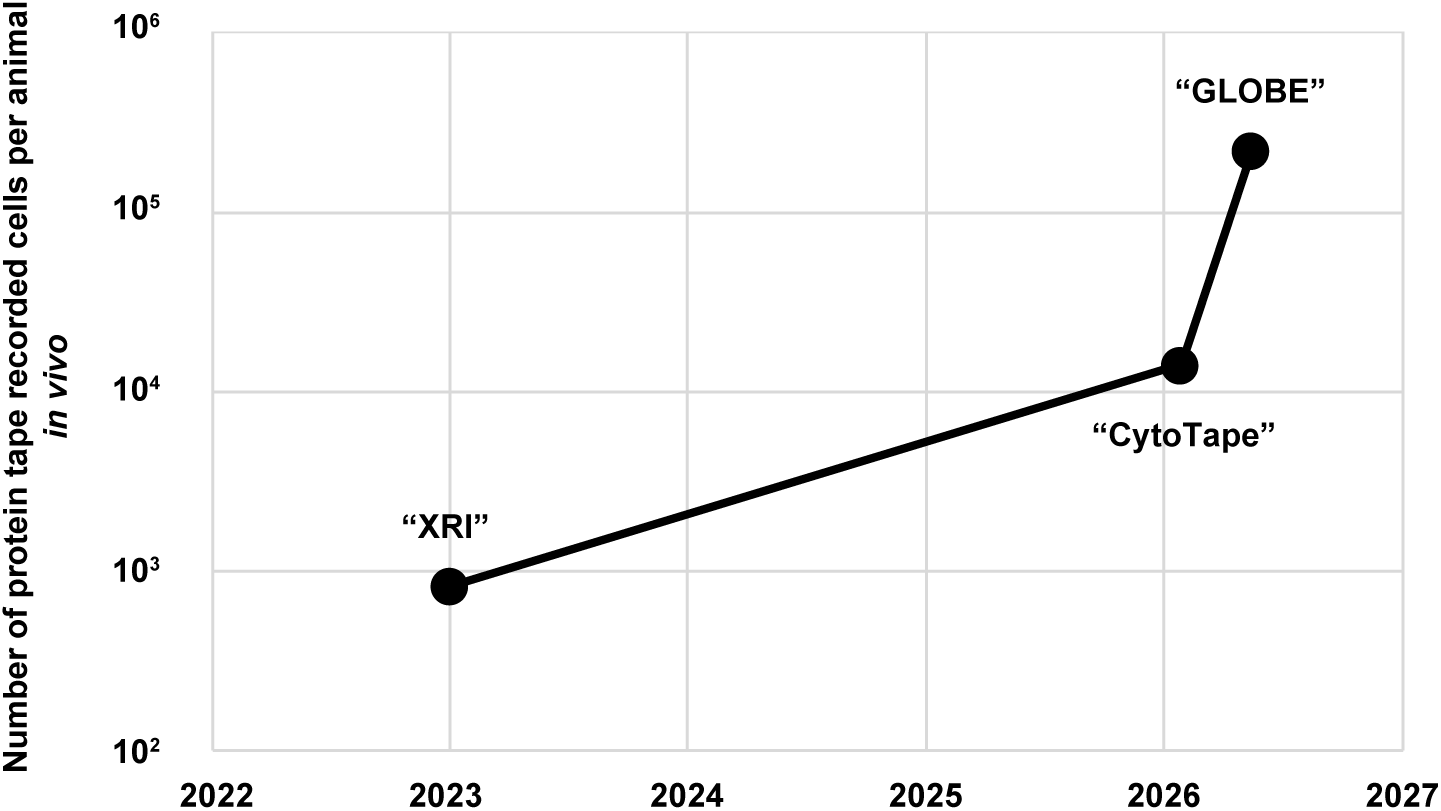
GLOBE scales up the throughput of simultaneous cellular recording *in vivo*. Throughput comparison of *in vivo* protein tape recording across successive systems. The “XRI” work recorded 835 neurons from one mouse^37^, whereas the “CytoTape” work increased the throughput to 14,123 neurons^38^. The GLOBE system in this work provided a recording throughput of up to 219,703 neurons from a single mouse brain, enabling simultaneous brain-wide single cell recording.

